# Macrophage-like vascular smooth muscle cells dominate early atherosclerosis and are inhibited by targeting iron regulation

**DOI:** 10.1101/2025.05.22.655434

**Authors:** Rijan Gurung, Chang Jie Mick Lee, Elisa A. Liehn, Siti Nurjanah, Ren Minqin, Jeroen A. Van Kan, Erielle Villanueva, Junedh M. Amrute, Shaun Loong, Matthew Ackers-Johnson, Francesco Ruberto, Shi Ling Ng, Yi Xuan Loo, Jiapeng Chu, Xiao Yun Lin, Konstantinos I. Karampinos, Theodoros Kofidis, Kory J. Lavine, Roshni R. Singaraja, Vitaly Sorokin, Roger S-Y. Foo

## Abstract

Vascular smooth muscle cells (VSMCs) contribute dynamically to atherosclerosis at all stages but the molecular drivers of their phenotypic switching, especially during early plaque development, and how they contribute to plaque progression remain unclear. We performed spatial transcriptomics on 12 human aortic tissues with and without atherosclerotic plaque. Macrophage-like SMCs were the predominant cell-type in the atheroma, displaying high iron storage and dysregulation, confirmed by spatial elemental mapping with nuclear microscopy. The combination of soluble iron and oxidized LDL promoted foamy macrophage-like VSMC cell state transition, while chelation inhibited this switching. *In vivo*, iron dysregulation induced neointimal thickening and macrophage-like switching in wire-injured *Ldlr^-/-^* mice, which was significantly reversed by ferrostatin-1, a ferroptosis inhibitor. These data show how targeting iron regulation modifies the macrophage-like VSMC cell state, and inhibits disease progression in atherogenesis.

## Introduction

Atherosclerotic cardiovascular disease is a leading cause of morbidity and mortality worldwide^1,2^. Build-up of an atheromatous plaque in the arterial wall underlies chronic, insidious atherosclerotic disease, while plaque rupture triggers acute thrombotic events, distal arterial occlusion, myocardial infarction (MI) or stroke^3–5^. Pathologically, the atherosclerotic plaque is initiated by an accumulation of low-density lipoprotein cholesterol (LDL-C) in the subendothelial space, followed by the formation of a lipid core containing inflammatory and vascular cells, and progressive accumulation of plaque debris and scar tissue^4,6^. Although macrophages were initially thought to be primary drivers of disease, recent evidence show instead that vascular smooth muscle cells (VSMCs) make up the majority of the plaque and drive neointimal thickening through their phenotypic modulation into at least 9 different cell states including VSMC that are macrophage-like, fibroblast-like and osteoblast-like^7–9^. Not only do these dynamic cells play a part in forming the fibrous cap over plaque lesions, they also contribute significantly to the overall plaque development at all stages^10^. Despite their key role in disease development and progression however, the molecular mechanisms behind VSMC cell state transitions, their progressive stages in plaque development, and whether precise cells states are suitable targets for disease therapy, remain to be fully elucidated.

Single-cell multi-omics and spatial transcriptomics offer a means to uncover new insights into precise cell types and disease mechanisms^10^. In the atherosclerotic plaque, this has so far shed light on the multiple players in plaque development, including endothelial cells, immune cells (especially macrophages), as well as “synthetic” VSMC and protective “fibromyocytes”^11–17^ in the context of plaque stability. However, a knowledge gap remains regarding cell state transitions in early plaque development.

## Results

### Spatial transcriptomic mapping of human atherosclerosis

In the present study, we conducted spatial transcriptomics on human vascular tissues obtained by tissue punch from the aortas of patients with widespread atherosclerotic disease (**Suppl Table 1**). Twelve plaque (“diseased”) and non-plaque (“control”) sections were analyzed by oil red O and Von Kossa staining to delineate the atheroma and confirm an absence of substantial calcification, respectively (**Fig 1A-B, Fig S1A-B**). Analysis using the Visium (10X Genomics) spatial transcriptomics platform (**Fig 1C**), revealed 18 clusters of vascular cells in diseased and control tissues combined (**Fig 1D-E, Fig S1C**). Eight clusters were identified as containing smooth muscle cells (SMCs), while others were fibroblasts, B cells, endothelial, myofibroblast-like, EndMT-like, NK/T cells, and macrophages. In particular, the SMC clusters showed diverse phenotypic cell states comprising of contractile SMCs (SMC-Con), SMCs transitioning towards a dedifferentiated phenotype (SMC-Trans), synthetic SMCs (SMC-Syn), fibroblast-like SMCs (SMC-Fibro) and macrophage-like SMCs (SMC-Mac) (**Suppl Table 2**). Notably, diseased atheroma sites contained almost exclusively SMC-Syn, SMC-Mac, and macrophage clusters (**Fig S1D-E**). Each cluster was characterized by highest expressed genes as shown in **Fig 1F** (**Fig S1F**). Because the Visium platform assesses transcriptomes as spots on tissue sections, we noted that the larger tissue sections and higher total spot counts in diseased compared to control sections were attributed to especially higher number and proportion of SMC-Trans, SMC-Fibro and SMC-Mac clusters (**Fig S1G**), again implying that in disease, plaque size expansion comprised of SMCs of these cells states.

**Fig. 1.**
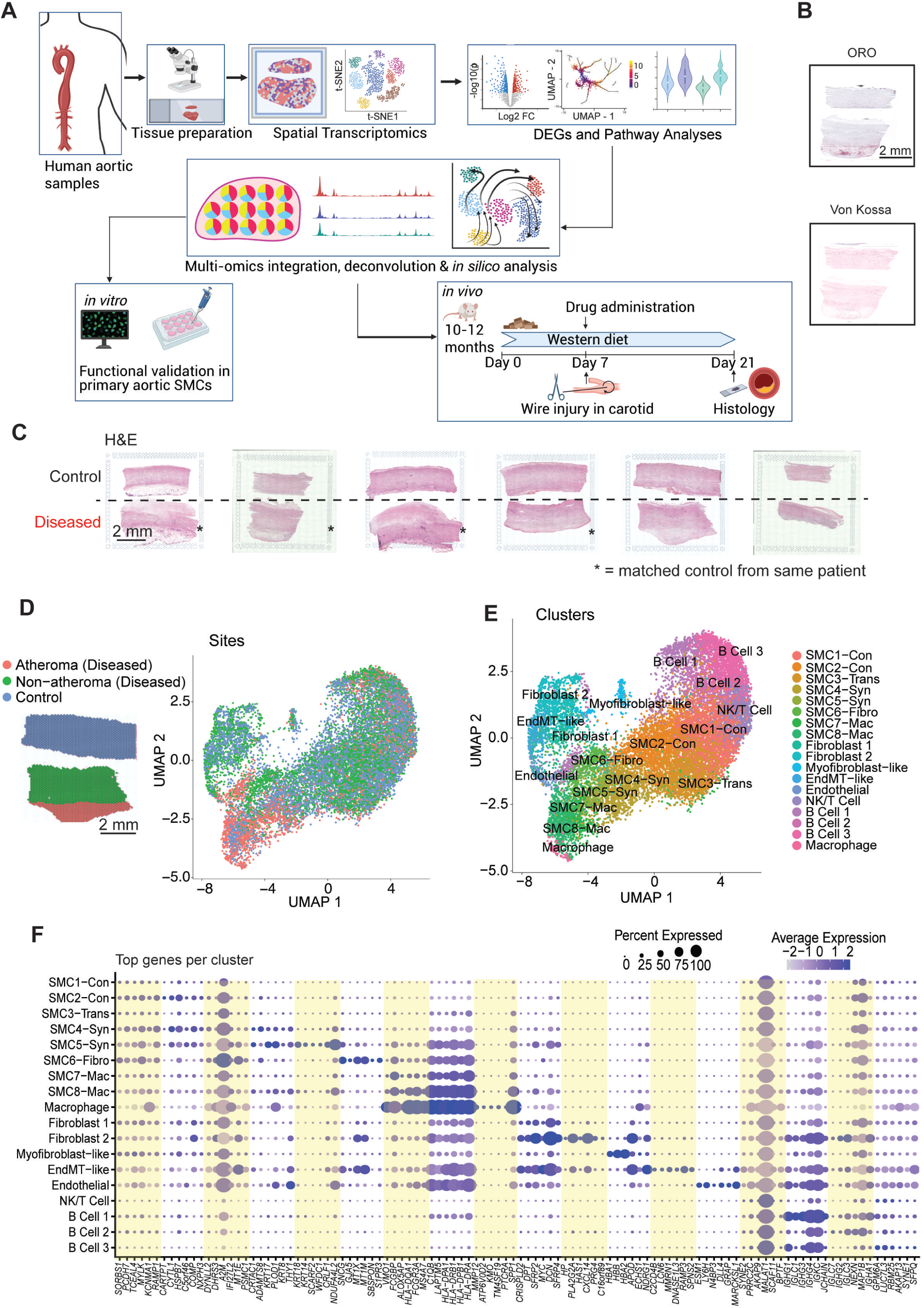
Study design, atherosclerotic plaque identification, and global clustering of human aortic tissues. **(A)** Study design. **(B)** Oil Red O and Von Kossa staining showing lipid deposition, but absent calcification, respectively. **(C)** H&E staining on Visium Gene Expression slides (n=6 Control without atheroma, n=6 Diseased with atheroma). **(D)** Spatial segmentation of control and diseased, atheroma and non-atheroma sites, and identification of site-specific spots located in an integrated UMAP embedding. **(E)** UMAP visualization of 18 clusters identified from all 12 tissues, in a total of 15,554 Visium spots. **(F)** Six highest expressed genes for each cluster are listed.

### Trajectory analysis of SMC cell state transitions in atherosclerosis

To characterize the SMC cell states, we used Monocle3^18^ pseudo-time analysis and delineated 2 distinct trajectories, starting from the contractile cluster and leading to either the macrophage-like cluster (Cell fate 1), or to the fibroblast-like cluster (Cell fate 2; **Fig 2A**). In succession, the analysis suggested that contractile SMC (SMC1-Con, SMC2-Con) dedifferentiated towards the transition state (SMC3-Trans) before arriving at the synthetic phenotype (SMC4-Syn, SMC5-Syn), and subsequently bifurcated towards either the macrophage-like (SMC8-Mac; Cell fate 1) or the fibroblast-like (SMC6-Fibro; Cell fate 2) cluster. Closer inspection revealed characteristic SMC genes: myosin heavy chain (*MYH11*), smooth muscle alpha actin (*ACTA2*) and transgelin (*TAGLN*) to be highest expressed in the contractile cluster, markedly decreased in the synthetic cluster, and lowest in the macrophage-like cluster (**Fig 2B**). In contrast, the synthetic gene osteopontin (*SPP1*) and macrophage markers galectin 3 (*LGALS3*) and *CD68* were lowest in the contractile cluster, elevated in the progressive switch to synthetic, and highest in the macrophage-like cluster. Unexpectedly, we noted a similar gene expression pattern in genes of the iron regulation pathway: ferritin light-chain (*FTL*) and heavy chain (*FTH1*), and transferrin receptor 1 (*TFRC*), transitioning through the synthetic to macrophage-like cell clusters.

**Fig. 2.**
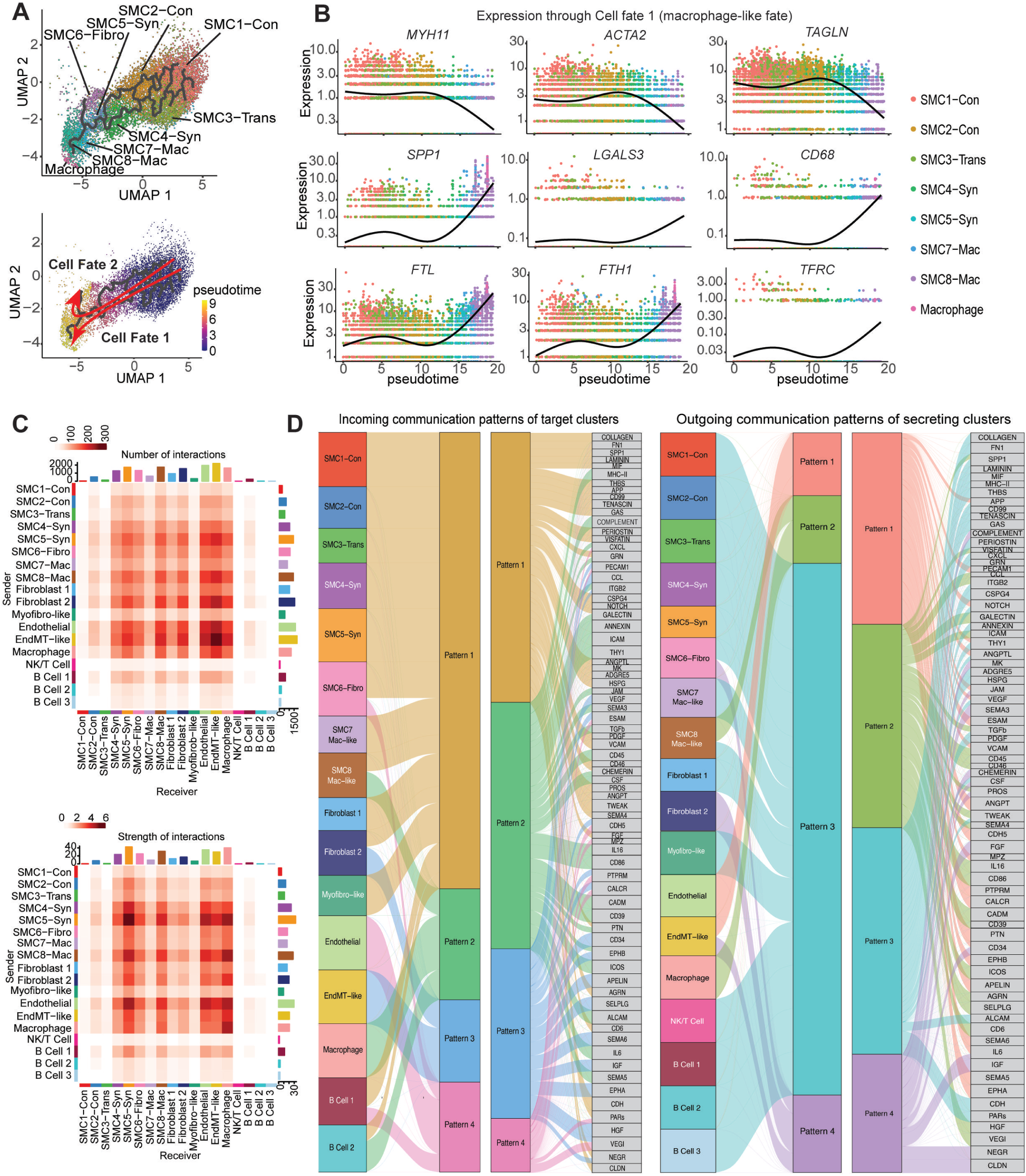
Trajectory analysis, cell-cell interaction, **(A)** UMAP visualization of 8 SMC clusters and Macrophage with trajectory analyses revealing two cell fates: Cell Fate 1 towards the macrophage-like cluster (SMC8-Mac), and Cell Fate 2 towards the fibroblast-like cluster (SMC6-Fibro). **(B)** Gene expression signatures of contractile markers (*MYH11*, *ACTA2, TAGLN*), macrophage-like markers (*SPP1*, *LGALS3*, *CD68*), and iron storage markers (*FTL*, *FTH1*, *TFRC*) through the Cell Fate 1 trajectory. **(C)** Cell-cell communication heatmap for Number of interactions (top) and Strength of interactions (bottom) between clusters as senders and receivers. **(D)** Riverplot depicting incoming and outgoing communication patterns of target and secreting clusters, behaving through key signaling pathways.

### Cell-cell communication in human atherosclerosis

Next, we carried out cell-cell interaction analyses in diseased samples using CellChat^19,20^. Compared to others, the SMC5-Syn, SMC8-Mac, Fibroblast 2, Endothelial, EndMT-like and Macrophage clusters had the largest number of ligand-to-receptor interactions when ranked for both senders and receivers (**Fig 2C**). In particular, the number of interactions from these clusters as senders to Endothelial, EndMT-like and Macrophage as receivers was highest, implying highly active cell-cell communication between these cell clusters in diseased sections. Similarly, the SMC5-Syn, SMC8-Mac and Endothelial clusters showed the highest strength of interactions with the same clusters, also implying highest abundance of ligand and receptor expression for cell-cell communication among them. Compared to control samples, cell-cell interactions of SMC6-Fibro, SMC7-Mac, SMC8-Mac, Fibroblast 2, and Endothelial among themselves and with Macrophages were markedly higher in diseased tissues (**Fig S2A-D**). Among all incoming and outgoing patterns of SMC cell-cell communication, our attention was drawn to the SMC8-Mac cluster because it shared both incoming and outgoing signaling patterns with other SMC (Pattern 1) and also with macrophages (Pattern 2; **Fig 2D**). Pattern 1 pathways were fibroproliferative and pro-remodeling (*FN1, SPP1, POSTN, NOTCH, PDGF, VCAM, ANGPTL, FGF*), while Pattern 2 pathways involved immune regulation, innate and adaptive immune signaling (MHC-II, complement, *CXCL*, galectin, *ICAM, THY1, CD45, IL16, CD86, IL6, SEMA3, SEMA5*).

### Spatial organization of cell clusters in human atherosclerosis

High assortativity scores in spatial analysis imply high likeliness for same cells to be located next to each other in the tissue spatially, while how likely pairs of different cells are next to each other is measured by the neighborhood enrichment score. Semla^21^ analysis showed that SMC8-Mac had markedly high assortativity implying that they had among the highest tendency to cluster together, and also with high neighborhood enrichment scores with macrophages and endothelial cells (**Fig S3A-D**). Other cells with highest assortativity scores included Macrophage, SMC4-Syn, Fibroblast 1 and SMC1-Con, implying that SMCs of the three major states in plaque development (contractile, synthetic and macrophage-like) tended to organize and cluster together spatially. In contrast, SMC8-Mac clusters were rarely found next to SMC-Con, SMC3-Trans or Fibroblast 1 clusters.

### Distinct transcriptomic signatures in vascular layers

In order to uncover specific gene programs that explain the putative cell states and their transitions, we analyzed deeper into the tissue sections and discerned distinct molecular signatures for each vascular tissue layer (**Fig 3A, Suppl Table 3**). Unsurprisingly, intima and atheroma sites in diseased tissues showed top gene signatures strongly implicating lipid accumulation (*APOC1*, *APOE, MSR1*), macrophage and foam cell activity (*CD68*, *LAPTM5*, *SPP1*, *CAPG*), immune activation (*HLA-DR*B1, *HLA-DRA*), inflammatory signaling (*CCL18*, *C1QB*), fibrosis (*POSTN*) and cell death (*KRT18*). Notably, top highly expressed ferritin light chain (*FTL*) and ceruloplasmin (*CP*) genes in the atheroma and media of diseased tissues, reflected significantly dysregulated iron pathways (**Fig 3A**). In contrast, the intima layer in control sections was characterized by homeostatic vascular maintenance (*CRLF1, PLAC9, MATN2*), protection against oxidative stress (*ADH1B*), cytoskeletal integrity and repair (*TMSB4X, S100A6*) and baseline immune regulation (*TFPI2*), reflecting preserved vascular integrity, limited oxidative stress or inflammation. Other vascular layers in the diseased tissues showed presence of inflammation and remodeling (*CP, FOS, IGKC, SFRP2, CXCL14*) as opposed to their control counterparts, which displayed elevated contractile (*TAGLN*) and anti-inflammatory genes (*APOD*, *SOCS3*). Top differentially expressed genes (DEGs) between diseased and control tissues were *SPP1*, *APOE*, *APOC1*, *LAPTM5*, *CD74*, *CD68* and *ITGB2* significantly elevated in disease (**Fig 3B, Fig S4A-B, Suppl Tables 4, 5**). Again, highly upregulated expression of ferritin, both light- (*FTL*) and heavy-chain (*FTH1*) in diseased sections characterized excessive iron regulated activity in the atherosclerotic lesions.

**Fig. 3.**
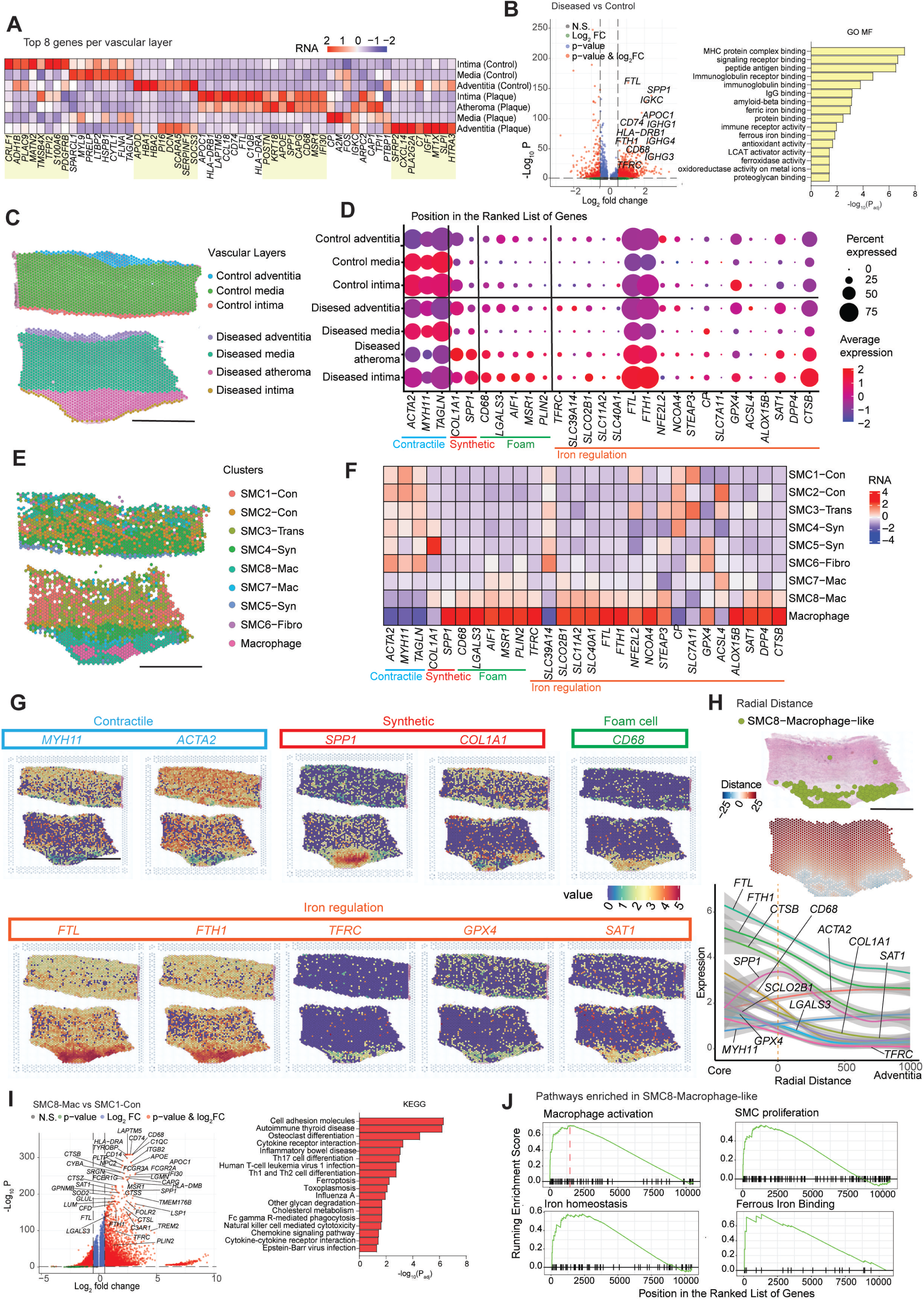
Characterization of vascular tissue layers, and altered iron regulatory gene expression. **(A)** Heatmap of top 8 expressed genes in each vascular layer for control (intima, media, adventitia) and diseased (intima, atheroma, media, adventitia) sections. **(B)** Volcano plot comparing diseased and control tissues, and gene ontology (GO) analysis. **(C-D)** Segmented vascular layers characterized according to SMC cell states (contractile, synthetic, foam) and iron regulatory gene expression. **(E-F)** SMC clusters were similarly characterized by cell states and iron regulatory gene expression. **(G)** Spatial map of gene expression for cell states and iron regulation. **(H)** Key gene expression for cell states and iron regulation measured across tissues by radial distance starting from the location of SMC8-Mac clusters in the atheroma (from atheroma core to adventitia). **(I)** Volcano plot comparing differential gene expression between SMC8-Mac and SMC1-Con, and KEGG enrichment analysis. **(J)** Gene set enrichment analysis (GSEA) reflecting pathways enriched in SMC8-Mac. Scale bar, 2mm.

### Iron dysregulation in atherosclerotic plaque

Dysregulated iron metabolism genes have been reported before in diseased atherosclerotic tissues^17,22,23^, but never analyzed in detail. To dissect this further, we studied gene expression in each layer of the tissue sections (**Fig 3C**). While contractile genes (*ACTA2*, *MYH11, TAGLN*) were enriched in the media of both diseased and control sections, and moderately expressed in control intima, they were downregulated in diseased atheroma and intima. On the other hand, the opposite was observed for synthetic genes *(SPP1, COL1A1*) and foam cell genes (*CD68*, *LGALS3*, *AIF1*, *MSR1*, *PLIN2*), which were expressed at low levels in control intima and media, and the media of diseased tissue, but markedly upregulated in the intima and atheroma of diseased tissue (**Fig 3D, Fig S4C**). Most strikingly, a similar pattern was seen for iron regulation genes, upregulated in diseased intima and atheroma, and expressed at low levels in corresponding control layers. Importantly, iron regulatory gene expression in each SMC cluster tracked also with phenotypic modulation (**Fig 3E-F**). SMC8-Mac showed downregulated contractile genes (*ACA2*, *MYH11, TAGLN*) compared to the other SMC clusters, and instead, upregulated synthetic (*COL1A1*, *SPP1*), foam cell (*CD68*, *LGALS3*, *AIF1*, *MSR1*, *PLIN2*), and iron regulation genes (*TFRC*, *SLCO2B1*, *SLC11A2*, *SLC40A1*, *FTL*, *FTH1*, *NFE2L2*, *NCOA4*, *STEAP3*, *ACSL4*, *SAT1*, *DPP4, CTSB*), mirroring the expression pattern of the macrophage cluster. The SMC7-Mac cluster contained a weaker expression signature compared to the more mature SMC8-Mac cluster, apart from elevated *ACS4*, implicating possible ferroptosis activity with increased availability of polyunsaturated fatty acids. Whereas *TFRC* appears as the primary iron uptake receptor upregulated in SMC8-Mac and macrophages, *SLC39A14* was primarily expressed for iron uptake in SMC-Fibro. Interestingly, ferroxidase *CP* was upregulated in SMC-Con, SMC-Trans and SMC4-Syn, but downregulated in SMC5-Syn, SMC-Fibro and SMC-Mac. We also observed site specific expression differences in the abundance of iron-regulation genes (*FTL*, *FTH1*, *TFRC*, *SLCO2B1*, *GPX4*, *SAT1*) that were upregulated in SMC8-Mac in the atheroma site, compared to non-atheroma sites in disease and control sections (**Fig 3G, S4D**). Moreover, across the radial distance of vascular thickness (**Fig 3H**), expression of contractile genes (*MYH11, ACTA2*) increased moving outwards from the atheromatous core to adventitia, whereas synthetic (*SPP1*, *COL1A1*), foam cell (*CD68*, *LGALS3*), and iron regulation genes (*FTL*, *TH1*, *TFRC, GPX4, SAT1*) increased inwards.

### Iron dysregulation in macrophage-like SMCs

Among all SMC cell states, the comparison between SMC8-Mac and SMC1-Con revealed distinct upregulation of foam cell markers (*MSR1*, *CD14*, *SPP1*, *TREM2*, *FOLR2*), markers of lipid metabolism (*APOC1, PLTP*), inflammation (*FCGR2A*, *TYROBP*, *GPNMB*, *FOLR2, LSP1, CFD*) and extracellular matrix (ECM) regulation (*LUM*) (**Fig 3I-J, Suppl Tables 6-8**), reflecting its cell state transformation. Specifically, iron uptake receptor (*TFRC*), ferritin (*FTL*, *FTH1*), genes related to ferroptosis (*SAT1*, *SOD2, GLUL*) and lysosomal activation (*LGMN*, *GPNMB*, *SLC14A3*, *CTSB*, *CTSS*, *CTSL*, *CTSZ*) were upregulated again in SMC8-Mac, whereas they were not in SMC-Syn **(Fig S5A-B, Suppl Tables 9-10)**. On the other hand, SMC8-Mac showed upregulated *TAGLN, ACTA2, MYL9* and ECM genes (*COL1A2, COL6A2*) compared to the Macrophage cluster, reflecting the characteristic of smooth muscle cells (**Fig S5C-D, Suppl Tables 11-12**).

### Spatial deconvolution of Visium spots confirms macrophage-like SMCs in atheroma

While the SMC8-Mac cluster co-expressed genes of both smooth muscle cells and foam cells, suggesting a macrophage-like SMC cell state transition, it was also possible that technically the Visium spot-based platform could not resolve if *bona fide* macrophages were closely proximate and juxtaposed to SMCs. We therefore used CARD^24^ to deconvolute the Visium spots in order to identify defined cell types. To this end, we integrated our spatial dataset with a published human atherosclerosis single-cell RNA-seq dataset^17^. By spatial deconvolution, the majority of atheroma in diseased sections were shown to indeed consist of macrophage-like SMCs (**Fig 4A**). Moreover, in the re-analysis of the dataset from Wirka *et al*, we could identify macrophage-like SMC (and other SMC states) marked by the gene signature that we had characterized (**Fig S6A-C**). Importantly, using two published datasets of *Myh11*^+^ lineage traced mice^17,22^, we also verified the diseased aortic SMC cell population that co-expressed foam cell marker *Cd68* and iron regulatory gene *Ftl1* (**Fig S6D-E**). In the deconvolution analysis, synthetic SMCs heavily lined the region between our tissue medial and atheroma layers, whereas contractile SMCs were located across layers of control tissues and medial layer of diseased tissues (**Fig 4B**). Synthetic SMCs scattered across control tissues likely reflected that these were not true healthy control aortic tissue, but were instead non-plaque regions from patients with widespread atherosclerosis.

**Fig. 4.**
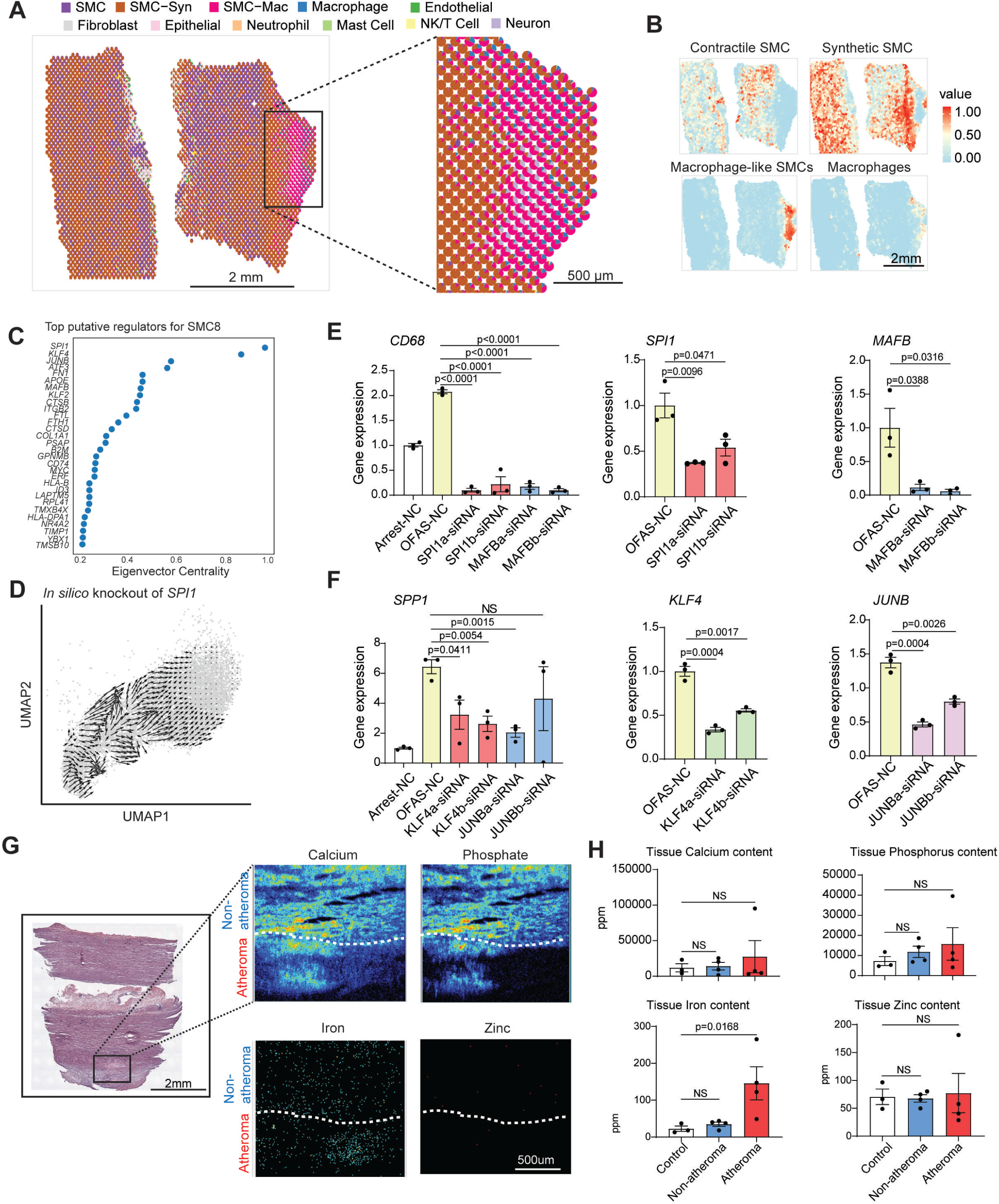
Deconvolution of Visium spots to validate SMC-Mac, top putative regulators of SMC cell state switching, and analysis of elemental iron in tissue sections. **(E)** A published human atherosclerosis single cell dataset^17^ was used as a reference to deconvolute our spatial data for each Visium spot. **(B)** Spatial enrichment of different SMC cell states in diseased and control tissues, visualized after deconvolution using the reference dataset^17^. **(C)** Top master regulators predicted by gene regulatory network analysis after integrating our data with a published single nuclear ATAC-seq dataset^26^. (D) *In silico* knockout of transcription factors (TFs) reversed SMC trajectory, away from macrophage-like, and towards the contractile state. **(E-F)** RT-qPCR analysis of *CD68* and *SPP1* in OFAS (50 µg/ml oxLDL and 5 µM ferrous ammonium sulfate/FAS) stimulated primary human aortic SMC after siRNA-knockdown of *SPI1*, *MAFB*, *KLF4* or *JUNB*. Data is presented as mean±S.E.M (n=3), one-way ANOVA test with Tukey’s multiple comparison test. **(G-H)** Spatial elemental mapping in tissue sections. H&E analysis (left) and nuclear microscopy (right) were performed and quantified for trace elements (iron, calcium, phosphorus, zinc) using Particle Induced X-ray emission (PIXE). Data is presented as mean±SEM, analyzed by Kruskall-Wallis test with Dunn’s multiple comparisons test.

### Putative TFs involved in SMC phenotypic switching

Next, to piece together potential molecular drivers for SMC phenotype switching, we used Cell Oracle^25^ in search for master regulators. We integrated our spatial transcriptomic dataset with a published single-nuclear ATAC-seq dataset^26^, and created a gene regulatory network (GRN). Using (a) *cis*-regulatory regions and gene expression data to infer transcription factor (TF)- target interactions, and (b) reference motif databases to identify motifs and map TFs, we quantified the strength of TF-target gene interactions in the SMC8-Mac cluster. These were ranked by degree centrality (signaling in/out/both), betweenness centrality and eigenvector centrality (**Fig 4C, S7A-B, Suppl Table 13**). Top TFs associated with macrophage-like switching that were also markedly upregulated in the SMC8-Mac cluster included *KLF4*, *ATF3*, *SPI1*, *KLF2*, *MYC*, *JUNB*, and *MAFB* (**Fig S7C**). By *in silico* knockout of each of *KLF4*, *ATF3*, *SPI1*, *JUNB*, and *MAFB*, we found strong reversal of the original cell fate trajectory, away from the macrophage-like phenotype and approaching closer towards a contractile cell state (**Fig 4D, S8A-B**).

To validate our findings thus far empirically, we knocked down each TF gene in aortic SMC *in vitro* using siRNAs and assessed for gene expression consequent to stimulation with a combination of oxLDL and soluble iron (ferrous ammonium sulfate, OFAS) (**Fig 4E-F**). SPI1 and MAFB are both ETS-family TFs previously implicated in macrophage differentiation. Knockdown of *SPI1* or *MAFB* suppressed *CD68* gene expression, while knockdown of *KLF4* or *JUNB* suppressed the expression of synthetic gene *SPP1*, suggesting that the 4 TFs may work cooperatively to promote pro-proliferative and macrophage-like SMC transitions. Most remarkably however, neither of these TFs affected iron regulatory gene expression following OFAS (**Fig S8C**). This revealed that alternatively, dysregulated iron metabolism could be acting instead as an upstream driver for SMC transitions, or running in parallel with the macrophage-like phenotypic switching.

### Spatial elemental mapping of human atherosclerosis

We therefore performed spatial elemental mapping using proton beam and particle-induced X-ray emission (PIXE) first to assess concentrations of the elements: calcium, copper, iron, phosphorus, and zinc in the same human aortic tissue sections (**Fig 4G**), in search for additional evidence of iron dysregulation. Indeed, in contrast to calcium, phosphorous and zinc, it was iron content that was significantly higher in diseased atheroma compared to control (**Fig 4H**). The high abundance of calcium and phosphorus, compared to iron, zinc and copper, in both diseased and control sections again likely reflected that all tissues were harvested from patients with widespread disease, and not true healthy controls.

### Macrophage-like cell fate with iron dysregulation *in vitro*

To test our emerging hypothesis for iron dysregulation in atherosclerosis, we stimulated cultured human aortic SMCs *in vitro* with IL-1β, PDGF-BB, carboxymethyl-lysine, and oxLDL (“IPCO”) and conducted RNA-sequencing to analyze their individual and combined effects on SMC phenotypic modulation (**Fig 5A-B**, **S9A-E, Suppl Tables 14-16**). Top differentially regulated genes were those for ECM remodeling (*MMP3*, *MMP10*, *MMP12*), inflammation (*IL1B*, *PTGS2*, *IL24*), metal homeostasis (*MT1G*), and pro-survival (*BCL2A1*). Gene ontology revealed processes of growth factor binding, chemokine binding, lipid binding, interleukin-1 receptor binding. Strikingly, haematopoietic cell markers indicated macrophage-like switching (**Fig 5B**). Enrichment for transition metal ion binding, oxidoreductase activity, ferroptosis, and mineral absorption corroborated data from our human spatial transcriptomes and elemental mapping.

**Fig. 5.**
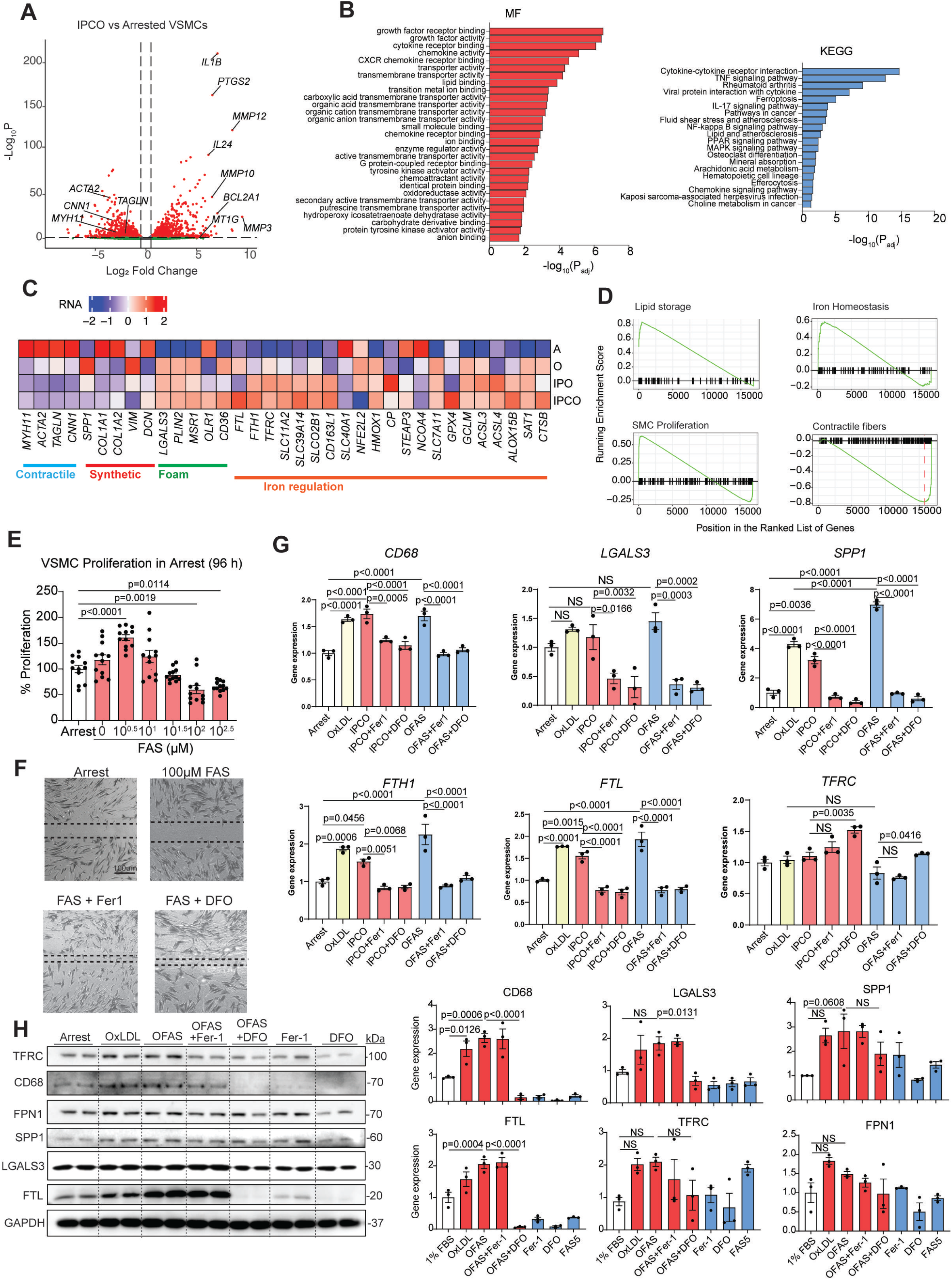
*In vitro* validation and functional analysis of the effect of iron metabolism on SMC phenotypic switching. **(A)** Volcano plot of differential gene expression and **(B)** GO analysis in IPCO-treated aortic SMC, compared to arrested control (IPCO; I=10 ng/ml IL-1β, P=10 ng/ml PDGF-BB, C=10 µM CML, O=50 µg/ml oxLDL). **(C)** Heatmap characterizing SMC transition towards a macrophage-like state after 48 h of stimulation with cocktail combinations. (A: arrested control, O=50 µg/ml oxLDL, IPO=10 ng/ml IL-1β+10 ng/ml PDGF-BB+50 µg/ml oxLDL, IPCO=10 ng/ml IL-1β+10 ng/ml PDGF-BB+10 µM CML+50 µg/ml oxLDL). **(D)** GSEA for enriched pathways following IPCO stimulation. **(E)** Proliferation of primary human aortic SMC after 96 h ferrous ammonium sulphate (FAS) administered in serum-free media. Data is presented as mean±S.E.M (n=12), one-way ANOVA with Dunnett’s multiple comparisons test. **(F)** Scratch assay measuring aortic SMC migration after treatment with 100 µM FAS, 5 µM Fer1 or 100 µM DFO for 8 h. Data is presented as mean±S.E.M.; one-way ANOVA with Dunnett’s multiple comparisons test. (**G**) RT-qPCR in SMC after combinations of O, IPCO, OFAS, with and without Fer-1 or DFO for 48 h (O=25µg/ml oxLDL, FAS=5µM). Data is presented as mean±S.E.M. (n=3), one-way ANOVA with Tukey’s multiple comparison test. **(H)** Western blot analysis for macrophage-related and iron regulatory proteins in SMC. Data is presented as mean±S.E.M. (n=3), one-way ANOVA with Tukey’s multiple comparison test.

oxLDL (“O”) alone pushed SMCs away from a contractile state by downregulating contractile genes (*MYH11*, *ACTA2*, *TAGLN*, *CNN1*) and strongly upregulating synthetic markers (*SPP1*, *VIM*), foam cell markers (*LGALS3*, *PLIN2*, *MSR1, CD36*) and iron regulation genes (*FTL, FTH1, TFRC, SLC11A2, SLC39A14, SLCO2B1, NFE2L2, HMOX1, SLC7A11, GCLM, ACSL3, SAT1, CTSB*) (**Fig 5C**). Adding pro-inflammatory and growth stimuli (IL1β, PDGF-BB) to oxLDL (“IPO”) pushed the cells from contractile and synthetic states, and even more to a macrophage-like state, with further elevation of iron storage (*FTH1, TFRC, SLC11A2, SLC39A14, SLCO2B1, CD163L1*) and ferroptosis genes (*ACSL4, ALOX15B*). Addition of the glycation end-product CML to mimic chronic atherosclerosis (“IPCO”), resulted in the strongest macrophage-like phenotype, with marked upregulation of genes for iron storage and sequestering, iron metabolism and ferroptosis. Gene set enrichment analyses of IPCO treatment included genes for lipid storage, iron homeostasis, and SMC proliferation, and poor enrichment for contractile fibers (**Fig 5D**). In order to establish a causal role for iron in the SMC phenotypic modulation response specifically, we administered soluble iron (FAS) to human aortic SMCs in culture. Despite a serum-free environment, FAS alone activated SMC proliferation in low concentrations (1-3 µM), although higher concentrations induced cell death (>100 µM) (**Fig 5E, Fig S10A-C**). Correspondingly, the iron chelator deferoxamine (DFO) inhibited serum-driven SMC proliferation. FAS also inhibited SMC migration (**Fig 5F, Fig S10D**), and co-administration of ferrostatin (Fer-1) or DFO with FAS reversed this effect. OxLDL, IPCO and OFAS all promoted phenotypic switching, shown by significant upregulation of *CD68*, *SPP1* and *PLIN2* (**Fig 5G, Fig S10E**). A moderate increase was seen for *LGALS3*, although this was not significant due to already high levels expressed in arrested cells. Similarly, levels of *FTL* and *FTH1* were significantly upregulated by the three treatments (**Fig 5G**). Co-administration of Fer-1 and DFO to both IPCO and OFAS treated cells significantly reduced *CD68*, *LGALS3* and *SPP1*, as well as *FTH1* and *FTL*. Although oxLDL, IPCO and OFAS did not alter expression of the iron importer *TFRC*, the addition of DFO significantly increased *TFRC* expression. At the protein level, OFAS significantly increased CD68 and FTL expression (**Fig 5H, Fig S10F**), whereas LGALS3, SPP1, TFRC and FPN1 increased after oxLDL or OFAS treatment. Co-administration of DFO with OFAS significantly reduced CD68, LGALS3, and FTL, and modestly reduced SPP1 and TFRC. FAS alone did not upregulate foam cell and iron related markers except for TFRC, suggesting that in this *in vitro* model, the dual presence of iron and oxLDL is needed to drive this phenotype.

Foam cell transition and lipid peroxidation of similarly stimulated human aortic SMC *in vitro* was also assessed using the Lipidspot and BODIPY C-11 dyes, respectively, with fluorescence microscopy. OxLDL treatment induced lipid uptake and peroxidation, which was further enhanced by FAS (OFAS, **Fig 6A-D**). Inhibition by Fer-1 or DFO significantly reduced lipid uptake and lipid peroxidation, consistent with the suppression of foam cell transition. As SMCs also undergo osteogenic transition in advanced atherosclerosis, we studied the effects of iron on human aortic SMCs cultured in calcification media (**Fig 6E-F**). Soluble iron, even at a high dose (300µM FAS), inhibited SMC calcification, whereas the ferroptosis inducer RSL3 significantly enhanced calcification, which was rescued by Fer1 or DFO. These implied that iron dysregulation, or even ferroptosis, appear to precede osteogenic transition.

**Fig. 6.**
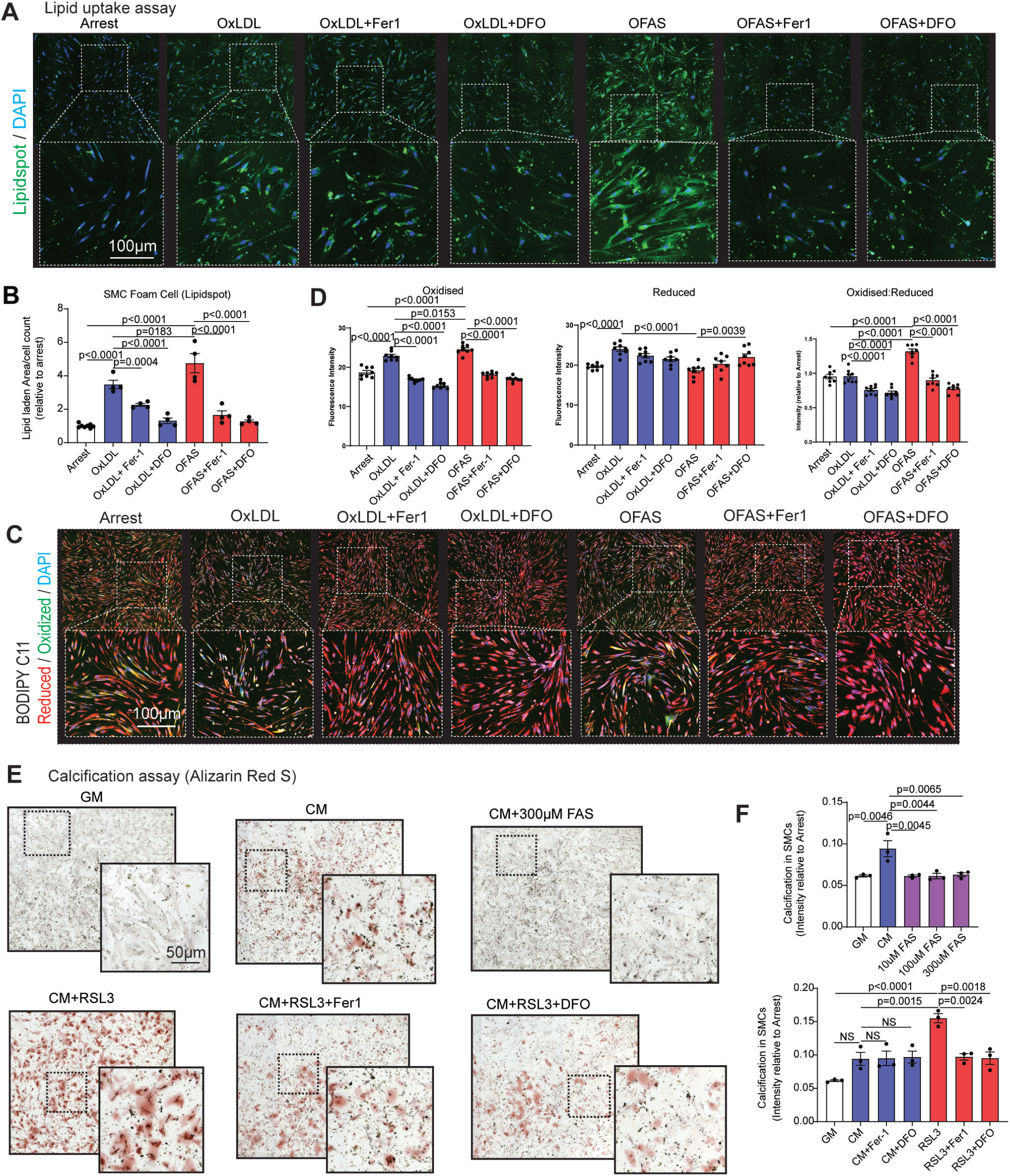
*In vitro* validation of the role of iron dysregulation in SMC phenotypic switching. **(A)** Lipid uptake indicated by LipidSpot (in green) and **(B)** quantified, in human aortic SMC after treatment as in Fig. 5. (Blue, nuclear DAPI). **(C)** Lipid peroxidation by BODIPY C-11 (red and green) and **(D)** quantified in aortic SMC after treatment as in Fig. 5. **(E)** Calcification of SMC (Alizarin Red S) and **(F)** quantified, after treatment with calcification medium (CM) compared to growth medium (GM), with combinations of FAS, 0.1 µM RSL3, 5 µM Fer1, and 100 µM DFO. Data is presented as mean±SEM (n=3), one-way ANOVA with Tukey’s multiple comparison test.

### Targeting iron regulation inhibits macrophage-like switching and plaque development ***in vivo***

In human atherosclerotic sections, total lesion area was approximately 8-fold larger compared to control sections (**Fig 7A-B, Fig S1A**). This was accompanied by a 3-fold increase in protein abundance of ferritin (FTL), CD68 and LGALS3 throughout the disease site. Hence, to causally validate the effect of iron dysregulation on SMC phenotypic modulation in atherosclerosis *in vivo*, we undertook wire-injury in *Ldlr*^-/-^ mouse carotid arteries to induce atherosclerosis, while also treating them with either the ferroptosis inducer (RSL3) or inhibitor (Fer-1) (**Fig 7C, Fig S11**). Wire-injury alone induced significant neointimal plaque formation, with abundant SMC accumulation as evident by αSMA staining. This was associated with increased CD68, LGALS3, FTL and TFRC protein abundance, which reflected macrophage-like switching, and increased iron uptake and storage (**Fig 7D**). Although RSL3 administration alone without wire-injury did not lead to substantial plaque buildup, RSL3 with wire-injury increased neointimal plaque size and TFRC levels, indicating that ferroptosis and iron uptake played an additive role to plaque expansion. In contrast, Fer-1 administration significantly reduced CD68, FTL and TFRC abundance, and markedly reduced neointimal plaque size, compared to wire-injury alone. Notably, the early transition marker LGALS3 expression was not reduced after Fer-1 treatment, showing that targeting iron regulation was effective in suppressing macrophage-like SMC switching more so than early synthetic transition. Moreover, even comparing between Fer-1- and RSL3-treated mice, there was a significant reduction in neointimal plaque size, and reduced expression of CD68, FTL and TFRC. Altogether, these findings explain how iron dysregulation can play a prominent role in macrophage-like SMC switching, and targeting this cell state altered the progression of atherosclerosis.

**Fig. 7.**
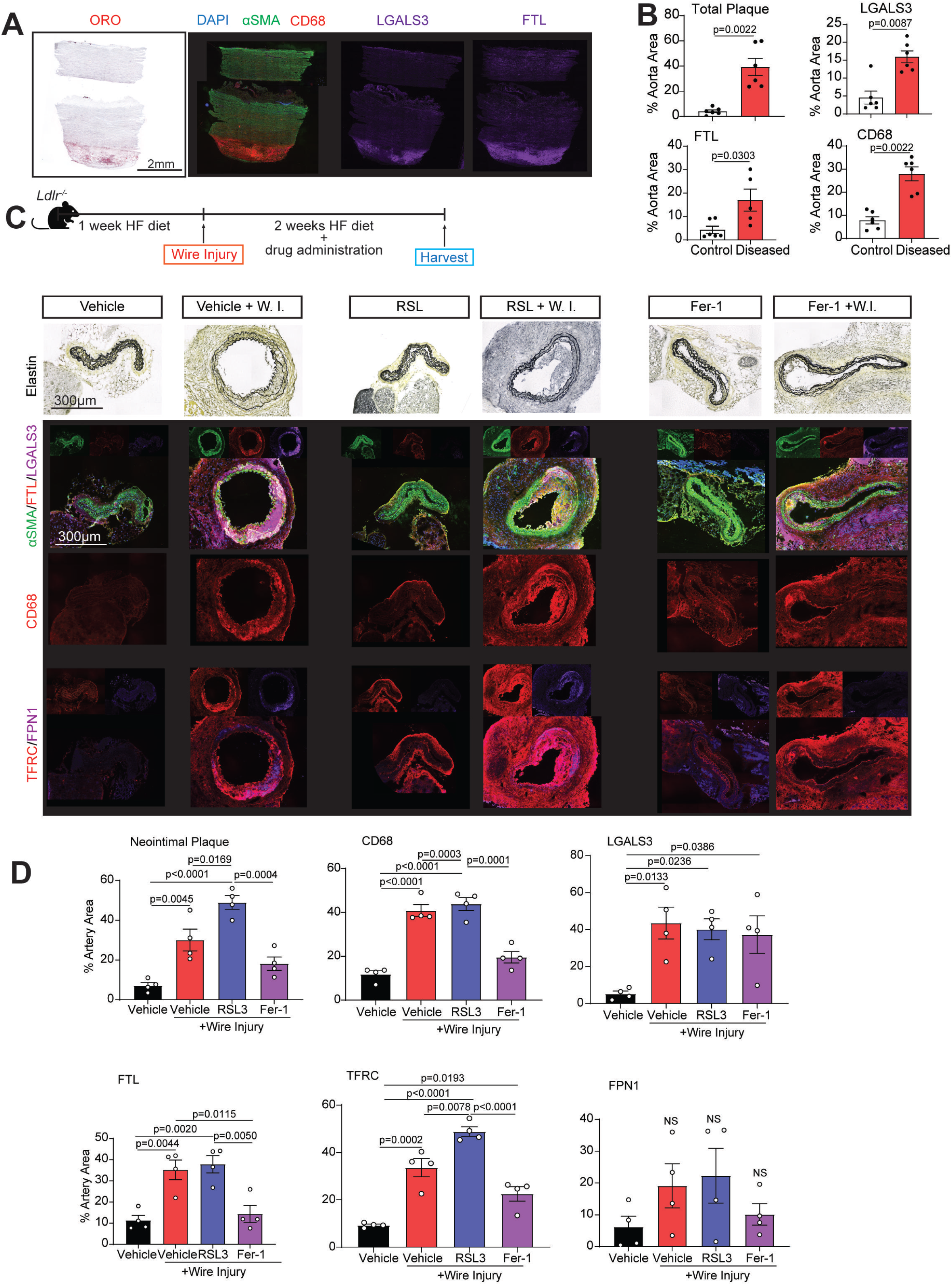
*In vivo* validation of the role of iron dysregulation in atherosclerotic plaque development. (A-B) Oil Red O and immunohistochemistry staining of human tissues, and quantification of total plaque area, macrophage and iron regulatory markers as a proportion of total aortic tissue area (LGALS3, purple; CD68, red; and FTL, purple) (n=6 per group). **(C-D)** Histological sections from wire-injured carotid arteries of *Ldlr*^-/-^ mice with either 10 mg/kg RSL3, 1 mg/kg Fer-1 or PBS control vehicle, showing elastin staining, αSMA (green), FTL (red), LGALS3 (purple), CD68 (red), TFRC (red), FPN1 (purple) and DAPI (blue). Neointimal hyperplasia and protein abundance relative to total artery area were quantified. Data is presented as mean±SEM (n=4), one-way ANOVA with Tukey’s multiple comparisons test.

## Discussion

By spatial analysis, we have mapped a transcriptomic landscape of the human atherosclerotic vascular wall. Vascular SMCs are dynamic cells central to atherosclerosis progression. They present in non-atheromatous regions as contractile or early synthetic cell states, transitioning towards a macrophage-like state only in areas at the atheroma. For the macrophage-like switching, we found it accompanied by a strong iron regulatory gene signature. This was seen in bulk and spatial transcriptomes, and further confirmed at the protein level in both mouse and human atherosclerotic plaques. *In vitro,* iron on its own promoted primary human VSMC synthetic transition, but added oxLDL challenge shifted it further to the macrophage-like state. Ferroptosis, on the other hand, promoted calcification in human VSMCs and increased neointimal growth in the *Ldlr*^-/-^ wire-injury mouse model. By integrating our spatial transcriptome with published chromatin accessibility data^26^ and linking transcription factor (TF) binding motifs to gene targets, we constructed GRNs and identified top putative master regulators of SMC phenotypic modulation.

From our analysis, macrophage-like SMCs comprised the majority of cells in the atheroma, likely because our tissues represented early, rather than late plaque disease. This is indeed the key novelty of our study because although other studies have used single-cell and spatial technologies to analyze atherosclerosis, their attention was focused on late plaque disease, plaque stability^17^ and rupture^16^, or other aspects such as acute and adaptive inflammation^11,27^. Sun *et al* spatially distinguished plaque rupture-prone regions as containing synthetic SMCs that express elevated MMP9, contributing to ECM remodeling, inflammation and fibrous cap thinning^28^. In terms of SMC phenotypic modulation, studies have highlighted the synthetic^29^, fibroblast-like^30,31^ and osteogenic phenotypic switching^23^, or analyzed non-SMC cell types such as macrophage^13,32^ or endothelial cell^14,15^, and the downstream myocardial response^33–35^. Data from Wirka *et al* revealed the transformation of SMCs into “fibromyocytes”, which was strongly driven by the transcription factor TCF21^17^. In that case, fibromyocytes provided a protective role through plaque stability. Alencar *et al* showed that contractile SMCs are driven by OCT4 towards a synthetic phenotype, and further modulated by KLF4 towards the osteogenic and potentially other states^16^. Similarly, Alsaigh *et al* identified VSMCs within the calcified atherosclerotic core as being matrix-secreting, osteogenic-like cells. Although phenotypic modulation had been studied, and SMC-like macrophages have been identified before in the plaque^30^, research has only emphasized the transition to synthetic intermediate cell states, and branching towards fibroblast-like or fibrochondrocyte-like cells^22,29,31,36^. Pathogenic foamy macrophages are described in the literature, but these were largely centered around cells of myeloid origin^13,32^. Gastanadui *et al* identified cells expressing the macrophage marker CD68 originating from endothelial, smooth muscle and myeloid lineages, which contributed to calcification and plaque instability^37^. Remarkably, the transcriptomic signatures of macrophage and macrophage-like SMCs in our study strongly resembled the foamy macrophages identified by Patterson *et al*^32^. Importantly, evidence from pulsed dual lineage tracing in atherosclerotic mice^38^ have supported the conclusion of SMC decreasing contractile markers and adopting a macrophage-like phenotype, instead of the counter possibility that myeloid cells acquire SMC gene expression, or indeed macrophages ingesting smooth muscle cells. Moreover, we have shown that targeting the iron regulatory pathway diverted the SMC state transition *in vitro*, and significantly reduced plaque size and the SMC plaque burden *in vivo*.

Our data of early plaque disease reveals at least 8 different SMC clusters, and 2 cell fate trajectories mapping transitions to either a macrophage-like or inflammatory fibroblast-like identity. Contractile cell clusters bore the hallmark signature of *ACTA2*, *MYH11* and *TAGLN* genes^39–41^, while transitioning SMCs downregulated these contractile genes and upregulated *SPP1*, a known synthetic marker^42^. Synthetic SMC clusters were additionally identified by expressing elevated levels of ECM-related genes *COL1A1*, *COL1A2*, *LUM* and *DCN*, a finding that is similar to the “fibromyocyte” identity previously described^17^. Macrophage-like cells instead had elevated *CD68*, *LGALS3*, *SPP1*^22^, whereas fibroblast-like SMCs had the ECM-remodeling signature with elevated *ADAMTS1*, *FOS*, *EGR* and *A2M*. We did not observe any osteogenic cell cluster, again reflecting early, rather than late-stage atherosclerosis in our tissues. By the use of full thickness aortic sections, we have successfully captured the full transcriptomic spectrum of different SMC states spatially in early disease transition.

As reported by others^16,22,36^, we also observed an intermediate cell state (ICS) that we call “synthetic” because these cells have not only upregulated genes typically observed during proliferation and ECM production, but also downregulated *MYH11*. Like others^16,22^, we also noted increase in *LGALS3* in synthetic and macrophage-like states, but in our data, this cell state was different from the SEM (stem cell, endothelial cell, monocyte) ICS described in past work^22^, as we did not find elevated *SOX9* and endothelial cell identity genes. Instead, our data is more consistent with the conclusion that contractile and fibroblast-like cells exist within a continuum^36^. More recently, it was shown that the ICS could de-differentiate further into either a fibromyocyte or fibrochondrocyte lineage, which also corroborates with our data^31^.

Interestingly, we found that *in vitro* macrophage-like switching showed downregulated ECM genes, while macrophage-like SMCs in human atheroma acquired an upregulated ECM signature, reflecting the more complex biology *in vivo* in human tissue. Nonetheless, increased *SPP1* expression was shared by both oxLDL-stimulated SMCs in culture and macrophage-like cells, and markedly reduced in synthetic and fibroblast-like SMCs, suggesting that elevated *SPP1* marks a synthetic phase driving towards the macrophage-like identity. Consistent with this, both *SPP1* and *POSTN* are highly abundant in late-stage atherosclerotic plaques^23^. SMC- macrophage clusters also highly expressed MHC-I and MHC-II molecules, suggesting potential interactions with CD4 and CD8 T cells^11,37^, and elevated levels of MMPs, as also noticed previously^28^.

Our study further reports for the first time that macrophage-like SMC transition is strongly governed by iron regulatory gene pathways, resolving a longstanding suspicion for their relationship in plaque development. Compared to other phenotypic states, macrophage-like SMCs bore significantly higher expression abundance of iron regulatory genes, which we also confirmed using another single-cell transcriptomic dataset of human atherosclerosis^17^ and two *Myh11*+ lineage traced atherosclerotic mouse models^17,22^. The same was also confirmed *in vitro* with primary human aortic SMCs stimulated by an atherogenic cocktail, even in the absence of serum. We show for the first time that soluble ferrous iron (FAS) enhanced macrophage-like switching, while iron chelator deferoxamine (DFO) inhibited this transition. Pertinently, by depleting ferrous iron, ferrostatin also inhibited the switch. In line with this, spatial elemental mapping confirmed increased total iron content in human atherosclerotic tissue, as also previously seen in a rabbit model of atherosclerosis^43^. *In vitro*, FAS and RSL3 suppressed SMC migration, suggesting that iron could also provide a stimulus for cell accumulation at the site of plaque burden to promote neointimal thickening. Similarly, iron enhanced foam cell transition *in vitr*o. As previously reported in a mouse model for chronic kidney disease^44^, ferroptosis promoted calcification in human aortic SMCs, which was abrogated by FAS administration. Taken together, our data now proposes a progression for how iron promotes macrophage-like phenotype in early-stage atherosclerosis and an osteogenic state in late-stage disease. *In vivo*, atherosclerotic plaques of wire-injured *Ldlr*^-/-^ mice composed of macrophage-like SMCs containing elevated ferritin, associated with increased neointimal thickening. Ferrostatin treatment markedly suppressed this, and reduced expression of *CD68*, but did not suppress *LGALS3*, potentially indicating that the cells remained in an intermediate poised state rather than reverting to fully contractile. These findings correspond with previous work using a ligation-induced atherosclerosis mouse model^45^, but now sheds light on the macrophage-like identity of VSMCs and the effect on iron-uptake and storage on this transition.

In conclusion, our work now provides a comprehensive spatial map of early atherosclerosis, showcasing the central and multi-dimensional role of vascular SMC plasticity, and how iron regulatory pathways govern plaque development. During disease progression, SMC identity transitions to an intermediate synthetic cell state before driving towards a macrophage-like state, making up the majority of cells in the atheroma. This precise phenotypic switching is not only influenced by lipoproteins but also by iron that is deposited in the lesion. Indeed, we show that targeting this cell state transition is alone sufficient to inhibit plaque disease progression. Taken together, this study now offers deep insight into early atherosclerosis, and points to new pathways for future therapeutic approaches.

## Acknowledgments

RSYF is funded by Individual Research Grants, and a Large Collaborative Grant from the National Medical Research Council (NMRC) of Singapore (MOH-001480-00) and MOE Academic Research Fund (AcRF) Tier 3 (MOE-000333-00). RRS is funded by grants from MOMMENTUM-CVD NMRC Centre Grant (CG21APR1008) and NMRC OF-IRG (MOH-001504-00). This work was partially supported by REDUCE-PAD National Research Fund Competitive Research Programme (CRP25-2020RS-0001). Primary human aortic SMCs were kindly gifted by Professor Shu Ye at Yong Loo Lin School of Medicine, National University of Singapore. We thank Ms. Nur Farhana Abdul Gani, Mr. Muhammad Haiman Bin Samad and Dr. Mario Ihsan for research coordinating support. We thank Dr. Caroline Chan and Dr. Matias Caldez for their assistance with the Visium platform setup. Study design schematics were created in BioRender.com. This publication is solely the responsibility of the authors.

## Author Contributions

RG and RSYF conceived the study. RG and RSYF drafted the manuscript with assistance from all authors. VS, XYL and TK collected human aortic tissue samples. RG and SL prepared human aortic tissues for spatial transcriptomics and performed 10x Visium cDNA construction for library preparations and sequencing. RG optimized and developed the *in vitro* atherogenic model and prepared all cDNA libraries for bulk RNA-sequencing. RG, CJML and EV performed all computational analysis. RG, SL and RM prepared human aortic tissues for spatial elemental mapping, which was subsequently performed and analyzed by RM and JAVK. RG, RRS, and EAL performed all in vivo experiments. RG, SL, and YL performed the histological and immunofluorescence data. RG, YL, JC and KIK performed all *in vitro* validation and functional experiments. RG, CJML, JMA, KJL and RSYF assisted with data interpretation of multi-omic data. All authors contributed to the experimental design, data analyses, interpretation and manuscript production. RSYF is responsible for all aspects of this manuscript, which includes project and experimental design, data analysis, and manuscript construction. All authors approved the final version of the manuscript.

