## Supplemental Materials and Methods for "Macrophage-like vascular smooth muscle cells dominate early atherosclerosis and are inhibited by targeting iron regulation"

**Ethics approval for human aortic samples**

Human atherosclerotic tissue sections (n=12 total) collected from 6 coronary artery patients undergoing coronary artery bypass surgery and control aortic tissues from 5 CAD patients and 1 thoracic aortic aneurysm patient were collected at the National University Hospital, Singapore. All procedures involving human patient samples were approved by the National Healthcare Group Domain Specific Review Board (reference number: 2009/00216 and 2013/00105) and informed consent was obtained from all participants. Data were collected and de-identified according to the National University Hospital Cardiovascular Tissue Bank (Tissue Bank Registration Number: NUH/2021-00047) protocol and ethical approval.

Samples were collected in Custodial HTK solution (Essential Pharmaceuticals, North Carolina, USA) and kept on ice before being processed for spatial transcriptomics analysis within 4 hours after resection. Aortic tissues were dissected, placed in O.C.T.-filled moulds and frozen in an isopentane-filled container placed in dry ice. The samples were then stored at -80^o^C until used experimentally.

**Spatial Transcriptomics processing and analysis**

Frozen aortic samples embedded in OCT (Sakura) were cryo-sectioned (Leica) at 14µm thickness and placed on pre-chilled Optimization slides (Visium, 10X Genomics, PN-1000193) and the optimal lysis time was determined following the Visium User Guide. The Nikon Eclipse TiE microscope and the NIS-elements software were used to capture images of fluorescently labeled cDNA. Spatial Gene Expression slides (Visium, 10X Genomics, P-1000187) were subsequently used for spatial transcriptomics following the Visium User Guides. Brightfield hematoxylin and eosin images were taken using a 10x objective on the EVOS M7000 microscope. Next generation sequencing libraries were prepared following the Visium user guide, and cDNA libraries were sequenced on a NovaSeq 6000 system (Illumina).

Visium fresh frozen data was aligned using Space Ranger (v2.1.0) and used for downstream analysis using Seurat (v5.2.1) and Semla (v1.0.0) in the R software (v4.3.0). Data was normalized using NormalizeData and ScaleData. nFeature_Spatial, nCount_Spatial and percent.mito were used to filter regressed data and remove low-quality spots. Spatial data from the different samples were integrated using Harmony (v1.2.1). FindVariableFeatures were used to identify highly variable and informative genes, followed by principal component analysis using FindNeighbors (reduction=“pca”, dims=1:20) used for dimensionality reduction. A k-nearest neighbours graph was constructed and clusters were identified using FindClusters (resolution=1.4) and visualized through UMAP generation (dims 1:20). Differentially expressed genes (DEGs) for each cluster were identified using FindAllMarkers. The resulting DEGs (adjusted p-value<0.05, top 100 Log2 fold-change) were submitted to Enrichr for pathway and functional enrichment analysis to aid in the annotation of each cluster. DimPlot was used to assess the clusters through UMAP visualization, while MapLabelsSummary was used to plot the spatial distribution of each cluster across the H&E stained images of aortic tissues. FeaturePlot was used to assess the expression of selected genes via UMAP visualization, while MapFeatures was used to assess gene expression spatially embedded over imaged tissues. FeatureViewer was used to select different tissues (control and diseased), sites (control, diseased non-atheroma, diseased atheroma), and layers (intima, atheroma, media, adventitia) for segmentation spatial analysis to identify region specific DEGs in the human tissues. Dotplot, VlnPlot and EnhancedVolcano (v1.18.0) with pCutoff=0.05 and Fccutoff=0.5 were used to visualize gene expression data through dot plots, violin plots and volcano plots, respectively. The Dittoseq package (version 1.12.2) was used to make stacked barplots with the dittobarplot() function, which revealed the number of spots per cluster and proportion of tissues represented by each cluster.

FindMarkers was used to identify DEGs between specific clusters and submitted to g:Profiler (adjusted p-value<0.05, top 200 genes ranked by adjusted p-value, followed by top 100 genes ranked by log 2fold-change) to obtain gene ontology and pathway enrichment analysis. The R packages MsigDB (v7.5.1, species = ‘Homosapiens’, category = ‘c5’) and clusterProfiler (v4.8.3) were used to perform gene set enrichment analysis (GSEA) and functional annotation of gene lists. Over-representation analysis (ORA) was performed using the enricher() function on upregulated and downregulated DEGs within each cluster of interest, such as the SMC8–Macrophage-like cluster. All DEGs for the cluster of interest were ranked by a GSEA metric, defined as the negative log10-transformed adjusted p-value multiplied by the sign (+1, 0 or -1) of the average log₂ fold-change. Genes were filtered to ensure uniqueness and absence of missing values, and the resulting ranked list was input into the GSEA() function with the MSigDB C5 gene sets. Enrichment results were exported as tables and visualized using the dotplot(), barplot(), and gseaplot() functions.

To investigate phenotypic transitions among smooth muscle cell (SMC) subpopulations, we performed pseudotime trajectory analysis using the Monocle3 package (v1.3.1). The Seurat object was converted into a Monocle cell_data_set object using the as.cell_data_set() function. To preserve the clustering and UMAP embeddings from our Seurat-based analysis, we manually assigned cluster identities and low-dimensional coordinates from Seurat into the Monocle object. All cells were designated to a single partition to enable unified trajectory inference. We constructed trajectories using learn_graph() and ordered cells along pseudotime using order_cells(). Visualization of the inferred trajectories and pseudotime distribution was performed using plot_cells(). SMC1-Con was selected as the root node as the biologically relevant early-stage cluster. To identify genes dynamically regulated along pseudotime, we performed a graph-based differential expression test using graph_test() with the principal graph, retaining genes with q-value < 0.05. Expression trends of selected lineage-associated genes (e.g., MYH11, TAGLN, CD68, FTL, FTH1) were visualized across pseudotime using plot_genes_in_pseudotime().

To perform cell-cell interaction analyses, we subsetted disease and control tissues from our spatial dataset and used CellChat (v1.6.1) and a pre-defined ligand-receptor pair database for human to obtain communication data and rank important interactions. To deconvolute the spatial data and determine the cellular composition of each spot, we integrated our data with a published single-cell transcriptomic dataset human atherosclerosis^1^ to identify cell types.

To infer and compare intercellular communication networks in spatial transcriptomic data from human aortic tissue, we used the R package CellChat (version 1.6.1). Seurat objects were first subsetted by tissue condition (control vs diseased) and CellChat objects were created using normalized gene expression matrices from the Spatial assay, alongside cell metadata. We used the human ligand–receptor interaction database provided by CellChat (CellChatDB.human) for downstream analysis. Following standard preprocessing, including the identification of overexpressed genes and interactions, we computed communication probabilities using a truncated mean approach with distance-based interaction modeling. We then inferred signaling pathways and aggregated the inferred networks to visualize both interaction number and strength through heatmaps using netVisual_heatmap. Specific signaling pathways (e.g., SPP1, PERIOSTIN, COLLAGEN) were visualized through heatmaps using netAnalysis_signalingRole_heatmap. To understand the roles of each cell type in signaling networks, we computed network centrality scores and clustered communication patterns using non-negative matrix factorization (NMF) to reveal distinct modes of signaling based on outgoing and incoming interactions. We visualized incoming and outgoing signaling roles through heatmaps using the netAnalysis_signalingRole_network() function, river plots using netAnalysis_river(), and scatterplots using netAnalysis_signalingRole_scatter().

To infer cell-type proportions at spatial transcriptomic spots in human aortic tissue, we used the CARD (Cell-type Assignment and Regulation Deconvolution) package (v1.1). To enable spatial deconvolution, we integrated our spatial transcriptomics dataset with a previously published single-cell RNA-sequencing (scRNA-seq) reference from Wirka et al. The single-cell data were re-annotated to reflect biologically meaningful cell-type identities described in the original study. We extracted the raw count matrix (sc_count) and metadata (sc_meta) from the single-cell data, and the spot-level count matrix (spatial_count) from our Visium spatial data. Spatial coordinates were imported using the LoadSpatialCoordinates function and filtered to match the barcodes present in the spatial count matrix. A CARD object was constructed using the createCARDObject() function, with parameters set to include all annotated cell types (ct.select = unique(sc_meta$cell.type)) and to retain genes and spots with at least one count (minCountGene = 1, minCountSpot = 1). The deconvolution was performed using the CARD_deconvolution() function, leveraging MuSiC (v1.0.0) for reference-based proportion estimation. To visualize spatial distributions of inferred cell-type proportions, we used the CARD.visualize.pie() and CARD.visualize.prop() functions, generating pie charts and cell-type-specific scatter plots of deconvolution results across spatial coordinates.

Spatial neighborhood statistics were performed using the Semla R package (v1.0.0). The diseased region was subsetted from the spatial dataset, and spatial proximity networks between Visium spots were computed using GetSpatialNetwork(), which establishes spatial edges based on adjacent spot locations. Clusters were visualized on the spatial network.

To assess spatial clustering of cell states beyond chance, label assortativity scores were calculated using RunLabelAssortativityTest() with 1,000 permutations. This metric evaluates whether cells of the same type tend to localize near each other in the spatial network. Results were visualized as scaled average degrees for each cell state. Next, to evaluate pairwise spatial interactions between different cell states, we ran the neighborhood enrichment test using RunNeighborhoodEnrichmentTest(), also with 1,000 permutations. This test computes the enrichment or depletion of neighboring label pairs, reporting results as Z-scores. A symmetric matrix of pairwise Z-scores was visualized as a heatmap. To highlight the most enriched and depleted interactions, the top 10 positively and negatively enriched label pairs were extracted and visualized in a bar plot. Radial distances from each spot to the closest spot in the SMC8-macrophage-like cluster were computed using the RadialDistance() function in SEMLA. Radial distances were visualized using MapFeatures, with square root transformation applied (r_dist_7_sqrt) to improve dynamic range and highlight local variation. We plotted smoothed expression curves of selected genes (e.g., *SPP1*, *ACTA2*, *CD68*) against radial distance using generalized additive models (GAMs). Dashed lines at zero distance indicated the approximate border of the reference cluster.

**Gene regulatory network construction and identification of key transcription factors**

To infer lineage-specific regulatory states during smooth muscle cell transitions, we integrated our spatial transcriptomic data with a transcription factor (TF)-target scaffold derived from single-nucleus chromatin accessibility profiles from a published dataset1. Gene regulatory network (GRN) inference was performed using CellOracle (v0.18.0) to construct predictive, cell-state-resolved regulatory networks. Processed scRNA-seq matrices, cluster annotations, and gene names exported from Seurat were imported into an AnnData object, which enabled downstream analysis with Python-based pipelines (Python, v3.13.3). The top 20 principal components and 4 nearest neighbors were used for initial graph construction and diffusion maps were computed with Scanpy (v1.12.1) to capture the manifold structure of the data. To refine neighborhood relations, the graph was subsequently recalculated using the diffusion map embeddings. An Oracle object was created by importing the processed AnnData and a pre-constructed TF-target scaffold generated from ATAC-seq peak-to-gene mappings1. Dimensionality reduction was performed, and the number of principal components retained was determined by identifying the inflection point of the cumulative explained variance curve, capped at 50 components. To mitigate the effects of technical noise and sparsity inherent in single-cell datasets, K-nearest neighbor (KNN) imputation was performed, where the number of neighbors (k) was set to 2.5% of the total cell count.

GRNs were then inferred for each annotated cluster using an elastic net regression model, balancing sparsity and stability (alpha = 10). Post-inference, links were filtered to retain only edges with p-values < 0.001, followed by selecting the top 2,000 most strongly weighted regulatory interactions for each cluster. Network statistics, including degree centrality, eigenvector centrality, and network entropy, were computed to capture key aspects of network topology. Comparative network analyses were performed to delineate transcriptional regulatory shifts between SMC and macrophage-like phenotypes. Centrality scores and network entropy values were visualized using scatterplots and boxplots. Dynamic regulatory programs associated with key transcription factors (e.g., SPI1, KLF4, MAFB, JUNB) were further interrogated through cartography plots, allowing visualization of their rewiring across cellular phenotypes. All GRNs, network scores, and ranked gene lists were exported for downstream integrative analyses. The full analysis workflow was performed using CellOracle (v0.18.0), Scanpy (v1.12.1), Seaborn (v0.12.1), and Matplotlib (v3.7.0).

***In silico* perturbation to assess and simulate cell state shifts**

The scaffold network was filtered to remove weak or unsupported interactions (links.filter_links()), and cluster-specific regulatory subnetworks were extracted (oracle.get_cluster_specific_TFdict_from_Links). GRN models were fit for simulation (oracle.fit_GRN_for_simulation) using Lasso regression with regularization strength alpha = 10.

To simulate the effect of transcription factor perturbation, we selected highest ranked TFs important for transitioning towards the SMC8 macrophage-like state as genes of interest (GOI). Simulated knockout was implemented by setting the expression of the GOI to zero (perturb_condition = {goi: 0.0}) and propagating signal shifts over three iterative steps (oracle.simulate_shift). Transition probabilities were estimated (oracle.estimate_transition_prob) using 200 nearest neighbors and a fully sampled cell population. Post-simulation, quiver plots were generated to visualize predicted identity shifts across the UMAP embedding (oracle.plot_quiver). To assess baseline noise, randomized simulations were performed (oracle.plot_quiver_random). For enhanced resolution, the embedding space was discretized into a 40x40 grid (oracle.calculate_p_mass) with smoothing (smooth=0.8), and a mass filter threshold (min_mass = 36) was applied to define reliable grid points (oracle.calculate_mass_filter). Simulation flow vectors were overlaid on both real and randomized grids for comparison.

Separately, pseudotime trajectories were inferred within the same embedding. Pseudotime values were discretized over the same grid structure (Gradient_calculator), and the developmental vector field was calculated using polynomial fitting (n_poly=7). The resulting pseudotime gradient was visualized independently and compared with simulated perturbation fields.

For direct quantitative comparison, an Oracle Development Module object (Oracle_development_module) was created, which enabled computation of the inner product between vectors from developmental flow and perturbation simulations, yielding a perturbation score (PS) for each grid point. Randomized control simulations were performed to establish significance thresholds. PS distributions were plotted across the entire manifold and visualized jointly with the perturbation vector field.

Focused analyses were conducted on specific lineages of interest. Cells belonging to predefined smooth muscle cell (SMC) and macrophage-like states were subsetted using cluster annotations (oracle.adata.obs['new_labels']). Separate Development Modules were created for SMC-derived macrophage lineages (Lineage_Mac) and SMC-to-fibroblast trajectories (Lineage_SMC). Perturbation scores were recalculated within these restricted populations to enhance lineage-specific signal detection. All analyses were performed using a Jupyter environment configured for full cell output (InteractiveShell.ast_node_interactivity='all').

**Single-cell transcriptomic analyses of human and *Myh11+* lineage-traced mouse atherosclerotic arteries**

We downloaded and re-analyzed data from two published datasets^1,2^. From the human atherosclerotic dataset, we identified a cluster within the macrophage population that substantially expressed contractile SMC markers *ACTA2* and *TAGLN*, which we considered to be of SMC origin and not myeloid. We subsequently used the Dotplot and VlnPlot functions to identify gene expression signatures of these subtypes according to smooth muscle cell states and iron regulation. We also subsetted smooth muscle cell lineage tdTmato+ cells in the WT group of the dataset from Wirka et al (*Nat Med,* 2019). Next, we used the AddModuleScore function from Seurat to create a gene set score and visualized using a RidgePlot. To characterize how our SMC-mac transcriptional signatures evolves in lineaged traced vascular SMCs in an *in vivo* murine model of atherosclerosis, we created a gene signature for our macrophage-like SMCs (*Cd68, Spp1, Lgals3, Pln2, Msr1, Cd36, Olr1, Trem2, Apoe, Laptm5, Cd74, Grn, Fabp5, Aif1, Lamp2,* and *Abca1*). Notably, we found that this Mac-SMC gene signature increases in vascular SMC lineage traced cells after 16 weeks of high fat diet compared to baseline. We also analyzed the Myh11-lineage traced (ZsGreen1^+^) SMC dataset from Pan et al (*Circ*, 2020) and identified macrophage-like SMCs enriched in mice after 26-week high fat diet. VlnPlot was used to assess macrophage-like and iron regulation gene signatures.

**Ion beam elemental tissue mapping**

Frozen aortic tissue sections of 15 µm thickness were collected on pioloform-coated nuclear microscopy target holders for elemental mapping and concentration analysis. A 2.1-MeV proton beam focused to a 1µm spot size was used to carry out elemental analyses. Scanning transmission ion microscopy (STIM), Rutherford back-scattering spectrometry (RBS), and proton-induced X-ray emission (PIXE) were simultaneously applied. A Silicon Drift Detector (SDD) X-ray detector placed at 90^o^ to the beam axis was used to detect X-rays of different elements simultaneously, as previously described^3^.

**Cell culture**

Human primary aortic smooth muscles were grown using human vascular smooth muscle basal medium (M231, Thermo Fisher Scientific) with smooth muscle growth supplement (SMGS, Thermo Fisher Scientific) in T-75 flasks before splitting using Accutase (STEMCELL Technologies) into appropriate cell culture plates for experiments. The experiments for aortic smooth muscle cells were conducted at passages 3-8 in Dulbecco’s Modified Eagle Medium with F-12 (DMEM/F-12; Thermo Fisher Scientific) supplemented with 0% or 2% fetal bovine serum (FBS; Sigma Aldrich). 1% penicillin streptomycin was added to all growth and arrest media.

**Bulk RNA-seq analysis**

Human VSMCs (Lonza) were stimulated with proinflammatory agents creating an atherosclerotic environment, *in vitro*. VSMCs were growth arrested in a serum free environment for 24 h, and subsequently stimulated using a cocktail (IPCO) of inflammatory mediators (10ng/ml interleukin-1β, 10ng/ml platelet derived growth factor-BB, 10µM Nε-Carboxymethyllysine, 50 ug/ml oxidated Low Density Lipoprotein), or further arrested over 48 hours. Total RNA was extracted using Trizol Reagent (Thermo Fisher Scientific) and Direct-zolTM RNA MiniPrep (Zymo Research) including DNase treatment using the manufacturer’s protocol.

RNA quality and quantity were assessed using the NanoVue Plus spectrophotometer (GE Healthcare) and the Agilent RNA 6000 Pico Kit (Agilent) which was run on an Agilent Technologies 2100 Bioanalyzer. Total RNA libraries were prepared using the Illumina TruSeq Stranded Total RNA Library Prep kit with Ribo-Zero Human/Mouse/Rat (Illumina, RS-122-2201) following the manufacturer’s instructions. cDNA libraries underwent QC using the Agilent DNA1000 kit (Agilent) before sequencing on an Illumina HiSeq4000 (~50 million paired-end reads per sample). Trim Galore was used for adaptor trimming, and reads were aligned to the human reference GrCh38/hg38 genome using STAR aligner with default parameters. Gene counts were computed using htseq-count, normalized to Counts per Million (CPM), and differential gene expression (DE) analysis was performed using EdgeR (version 4.0.0; false discovery rate-adjusted P value <0.05, Log fold-change>1). Figures were generated with R (Version 4.3.3).

**Transfection with siRNA *in vitro***

VSMCs were seeded into 24-well plate and grown to confluence, and were transfected for 24 hours with either SPI1, MAFB, JUNB, KLF4 siRNAs or negative control siRNA (Supplementary Table S17) at 25nM using Lipofectamine RNAiMAX Reagent (Thermo Fisher) in accordance with the manufacturer's protocol. The media was changed to arrest medium (0% FBS in DMEM/F12) for another 24 hours, after which the cells were treated with 25ug/ml oxLDL and 5uM ferrous ammonium sulfate (FAS).

**Iron detection *in vitro***

FerroOrange (Dojindo) was used to stain for intracellular Fe^2+^ in VSMCs. VSMCs were growth arrested for 24 hours before treatment with ferrous ammonium sulfate (FAS; Sigma Aldridge), 10µM ferrous ammonium sulfate (FAS), 5 μM ferrostatin-1 or 100 μM deferoxamine (DFO). VSMCs were treated with 1 μM of FerroOrange prepared in DMEM/F12 for 30 min at 37^o^C and washed with PBS. Staining was assessed by Nikon Eclipse TiE microscope and the NIS-elements software, and quantification was performed using ImageJ software.

**VSMC proliferation assay**

To assess proliferation, cells were plated in 48 well plates at 2 x 10^4^ cells/mL density and growth arrested for 24 hours prior to treatment. Cells were treated with 30 µM FAS, 5 µM Fer-1, and 100 µM DFO in DMEM/F12 with 2% FBS and incubated for 96 hours at 37^o^C. The cells were fixed with 4% (w/v) PFA for 15 min, permeabilized for 5 min using 0.1% triton X-100 and stained with 1 μg/ml DAPI for 5 minutes at room temperature. Cell proliferation was measured by counting total cell number provided by the nuclear stain using ImageJ.

**VSMC migration assay**

VSMCs were seeded in 24 well plates and grown to confluence, after which they were growth arrested for 24 hours and scratch wound assays were conducted. A scratch was created using a p200 pipette tip through the center of each well, which were then washed with PBS and treated in the appropriate treatments in serum free DMEM/F-12. VSMCs were incubated at 37^o^C for 8 hours or 15 hours, fixed in 4% (w/v) PFA for 10 min. The cells were stained with 0.05% Crystal violet for 30 minutes in room temperature and washed twice with tap water. Images were taken at times 0 and 8 hours or 15 hours using an EVOS M7000 microscope and migration was calculated as a percentage of total area at 0 hours using ImageJ.

**VSMC viability assay**

VSMCs were seeded into a 96-well plate and grown until 90% confluence. They were subsequently growth arrested for 24 hours and treated with 100 µM and 300 µM ferrous ammonium sulfate (FAS), 5uM ferrostatin-1 (Fer-1), 100 μM DFO, and 3µM and 10 µM RSL3 for 72 h in serum-free media (DMEM/F-12). 10uL of the Cell Counting Kit-8 (CCK-8) reagent (Abcam) was added to every well and the VSMCs were incubated for 2 h at 37^o^C. The optical density (OD) at 450 nm was determined using a plate-reader. Cell viability was expressed as the percentage of the OD of the experimental group compared to the control group.

**Real time quantitative PCR analysis**

Total RNA was harvested using TRIzol (Thermo Fisher Scientific) and RNA was extracted according to the manufacturer’s protocol. cDNA was synthesized using oligo dT and random primers provided with the HiScript III cDNA Synthesis Kit (Vazyme). A total of 200ng of total RNA was reverse transcribed. 4ng of cDNA was combined with 0.5µM of qPCR primers and 5µL of Low Rox PerfeCTa SYBR Green Fast Mix (Quanta Biosciences). RT-qPCR was performed on a QuantStudio 5 real-time PCR system. The primers and sequences utilized in this study are detailed in Supplementary Table S18.

**Western blot analysis**

VSMCs were grown to confluence, growth arrested for 24 h and treated in media containing 2% FBS with 25µg/ml oxLDL, 5 µM FAS, OFAS (oxLDL+FAS), 5 µM Fer-1 and/or 100 µM DFO for 48 h at 37^o^C. Cells were washed once with cold PBS and lysed using RIPA lysis buffer (Tris, NaCl, NP-40, sodium deoxycholate and 0.1% SDS) supplemented with protease inhibitor cocktail (Thermo Fisher Scientific). Cell lysates were centrifuged at 20,000g for 20 minutes to remove cell debris. Proteins were measured using the Pierce BSA Protein Assay Kit (Thermo Fisher Scientific) as per manufacturer’s protocol. Laemelli buffer with 10% β-mercaptoethanol was added to each sample and the mixture was heated to 90^o^C for 5 min. The Western blot was run using 12% gels at 80V for 2 h, and transferred to a PVDF membrane at 0.35A at 100V, which was incubated with primary and secondary antibodies in 5% milk in TBST. The immunoblots were developed with chemiluminescent substrate (Thermo Fisher Scientific) and visualized with BioRad ChemiDoc Imaging System. The blots were analyzed using ImageJ and normalized to GAPDH. Primary and secondary antibodies and reagents are listed in Supplementary Table S17.

**Lipid loading assays**

VSMCs were grown to 80% confluency in M231 medium supplemented with SMGS prior to experimentation. The media was changed to DMEM/F-12 with 2% FBS for 24 hours followed by treatment with oxLDL. Lipid accumulation was measured by administering 25μM oxLDL (Thermo Fisher Scientific) for 48 h to the VSMCs and adding 1 μM Lipidspot (Biotium) for 30 min at 37^o^C in the dark. 2 μM BODIPY-C11 was used to assess lipid peroxidation. The cells were washed with phosphate buffered saline (PBS) and fixed in 4% (w/v) PFA for 15 min and incubated with 1 μg/ml Hoescht (Sigma) for 10 minutes to stain for the nucleus. Images were captured by the EVOS M7000 microscope. Quantification of lipid loading was performed using the ImageJ software.

**VSMC calcification assay**

VSMCs were grown to 70% confluence in 24 well plates and administered with treatment conditions that were prepared in calcification media consisting of DMEM/F-12 supplemented with 10% FBS, 1% penicillin streptomycin, 5mM β-glycerophosphate (Sigma), 3 mM CaCl^2+^, 50 μM L-AA, and 100 nM dexamethasone. VSMCs treated with growth medium (DMEM/F-12 with 10% FBS, 1% penicillin streptomycin), regular calcification medium with/without FAS (10µM, 100µM, 300µM), 0.1µM RSL3, 5µM Fer-1 or 100µM DFO were incubated for 7 days. The cells were fixed in 4% (w/v) PFA for 15 min and washed once with ddH_2_O. To assess for calcification, 2% Alizarin Red S (ARS) solution prepared in ddH_2_O and adjusted to pH4.2 was applied for 30 minutes at room temperature. The cells were washed twice with ddH_2_O prior to taking images using the EVOS M7000 microscopy system. To quantify levels of calcification, 10% citric acid was used to dissolve the ARS at R.T. for 30 minutes. Samples were transferred to a 96-well plate reader and read at 405nm.

**Animals and treatment regimes**

All mouse experiments were approved by the Biomedical Sciences Institute Singapore Institutional Animal Care Committee at A*STAR (166165). Male, 10 to 12 week *Ldlr*-/- mice (C57BL/6J background) were fed lipid-rich Western-Type Diet 1 week before and 2 weeks after wire injury. Mice were anesthetized (100 mg/kg ketamine hydrochloride, 10 mg/kg xylazine i.p.) and subjected to endothelial denudation of the left common carotid artery using a flexible 0.36 guide wire through a transverse arteriotomy of the external carotid artery, as previously described^4^. Mice were randomly divided into three treatment groups (PBS, RSL3, and Ferrostatin-1 (Fer-1) treatment) with two arms each (non-injury and wire injury). 10 mg/kg RSL3 (Selleck Chemicals) and 1mg/kg Ferrostatin-1 (MedChemExpress) were administered via intra-peritoneal injection once a day beginning on the day of wire injury for the duration of the two weeks. Mice were then anesthetized (100 mg/kg ketamine, 10 mg/kg xylazine, i.p.) and carotid arteries were excised, embedded in O.C.T. in cryomolds and fresh frozen at -80^o^C.

**Histological analysis and immunofluorescence staining**

Aortic sections were sectioned at 10µm thickness (Leica cryostat) and re-hydrated for hematoxylin and eosin (H&E) staining and Von Kossa staining, according to the manufacturers’ protocol, followed by imaging using the EVOS M7000 microscopy system. For immunofluorescence staining, aortic sections were sectioned at 5µm thickness and re-hydrated, fixed in 4% paraformaldehyde and washed before permeabilization using 0.5% Triton X-100/DPBS. The sections were blocked in DPBS containing 5% donkey serum and 0.1% Triton X-100 at R.T. 1 hour followed by incubation in primary antibody overnight at 4^o^C. Secondary antibodies were applied at R.T. for 1 h the next day and sections were incubated in 1% Sudan black B in 70% ethanol at R.T. for 7 min to reduce the autofluorescence prior to final mounting. Images were captured by the EVOS M7000 microscope. Quantification of neointimal plaque size and protein abundance relative to total artery area were performed using the ImageJ software. Primary and secondary antibodies and reagents are listed in Supplementary Table S17.

**Statistics**

As sample size was governed by tissue availability, no sample size calculations were performed. No samples were excluded. For human samples, all collections were randomized across age, sex and race. For *in vivo* studies, male mice were randomized to control and treatment groups, and all histological and plaque quantification was performed blinded. All spatial and single cell transcriptomic, and single-nuclear epigenetic analyses were performed using un-biased techniques with the investigators being blinded when performing initial analyses. Data are presented as mean ± SEM. Shapiro-Wilk test was used to examine normality in data. For normally distributed data, two-tailed, unpaired Student’s t test was performed for comparisons between two groups, while one-way ANOVA followed by either Tukey’s or Dunnett’s multiple comparison test was used to compare more than two groups. For non-normally distributed data, Mann-Whitney U test was performed for comparisons between two groups, while Kruskal-Wallis test with multiple comparison test was used to compare more than two groups. All tests were performed using GraphPad Prism 9. P<0.05 was considered significant.

**Data availability**

Raw and processed sequencing files can be accessed on the Gene Expression Omnibus super series (…)

**Code availability**

Scripts used for analysing the data in this manuscript can be found at (https://github.com/rijgrg)

**Fig. S1. Human aortic control and diseased tissues, overview of clusters and Visium spot assignment. (A)** Oil Red O staining to visualize neutral lipids in human aortic tissues: control (top) and diseased sections (bottom). **(B)** Von Kossa staining showing absence of prominent calcification. **(C)** Spatial distribution of clusters in 12 tissues (6 control and 6 diseased). **(D)** Spot assignment across vascular layers used for segmentation analysis. Scale bar, 2 mm. **(E)** Spot distribution of clusters in control and diseased sections. **(F)** Characterization of 18 clusters using known gene markers. **(G)** Spot count and proportion of clusters in tissues captured on the Visium platform. Red arrows point to larger number and proportion of spot clusters SMC5-Syn, SMC6-Fibro and SMC8-Mac in diseased tissue.

**Fig. S2. Cell-cell interaction analyzed by CellChat. (A)** Net differential number of interactions and strength of interactions between clusters. Red indicates higher levels of interaction in diseased tissues, blue indicates higher levels of interaction in control tissues. **(B)** Ranking of incoming and outgoing interaction strengths for each cluster in diseased tissues. **(C)** Ranking of outgoing and incoming signaling patterns of each cluster in diseased tissues. **(D)** Heatmap of top signaling pathway networks relevant specifically for SMC8-Mac. The strength of signaling via the pathway network with every cluster as either sender, receiver, mediator or influencer to each of the other clusters is shown.

**Fig. S3. Spatial statistics, DEG analyses of tissues, and DEG between SMC5-Syn and SMC8-Mac clusters. (A)** To obtain statistics for spatial relationships, Visium spots were converted to nodes in a spatial graph using spot coordinates, forming a K-nearest neighbor (KNN) graph. Scale bar, 500 µm. **(B)** Assortativity scores measuring the degree of spatial clustering between same spot identity, where high scores represent highly non-random clustering, and low scores represent spatial mixing or dispersion of the same spots. **(C)** Heatmap of Neighborhood enrichment scores measuring degree of spatial clustering between different spot pairs, and **(D)** ranked via Z-score.

**Fig. S4. (A)** Spot assignment in control and diseased tissues. **(B)** Enriched pathways from differential expressed genes (DEGs) between diseased vs control tissues using Gene Ontology Biological Processes (GOBP) and Wiki Pathways (WP). **(C)** Spatial map of contractile, synthetic, foam, and iron metabolism gene expression in control and diseased tissues. **(D)** Violin plots of iron regulatory genes (*FTL*, *FTH1*, *TFRC, SLCO2B1, SAT1*, *GPX4*) in control tissues, and diseased non-atheroma and atheroma sites. The SMC8-Mac cluster is highlighted in yellow box.

**Fig. S5. (A)** Volcano plot and **(B)** GO Biological Processes (BP) and GO Molecular Functions (MF) of DEGs in SMC5-Syn compared to SMC1-Con. **(C)** Volcano plot and **(D)** GO BP and GO Cellular Component (CC) of DEGs in SMC8-Mac compared to Macrophage.

**Fig. S6. Re-analysis of a published single-cell human coronary artery disease (CAD) dataset, and 2 mouse *Myh11*^+^ lineage traced atherosclerosis datasets, revealing *bona fide* macrophage-like SMCs showing upregulated ferritin and iron dysregulation. (A)** UMAP and feature plots showing identification of macrophage-like SMCs in a published human CAD dataset by Wirka et al^17^. What was labelled as Macrophages in the published figure (left UMAP) in fact contained elevated expression of SMC genes *ACTA2* and *TAGLN* (2 plots on the right, selected in orange). Cells in this cluster co-expressed macrophage-like genes *CD68*, *LGALS3*, *PLIN2*, *TREM2* and *FABP5*. Re-analysis of the Macrophage cluster showed the presence of what we have re-labelled as SMC5-Mac (lower panel), which we believe are macrophage-like SMC, closely sharing a gene signature profile to SMC8-Mac in our dataset. **(B)** Dot plot summarizing gene expression patterns of all cells in the human CAD dataset, identifying upregulated genes involved in iron regulation in SMC5-Mac cells. **(C)** Violin plot showing macrophage-like SMCs expressing reduced levels of contractile gene *ACTA2* and markedly elevated macrophage marker *CD68* and light-chain ferritin (*FTL*), resembling the macrophage cluster. **(D)** From one *Myh11*^+^ lineage traced mouse dataset^17^, upregulated pseudobulk recovery signature ridge plot split across the 3 time-points of high fat diet, show macrophage-like SMCs appearing enriched in 16-week high fat diet treated mice. **(E)** Violin plot of a second dataset of *Myh11*^+^ lineage traced SMCs from mice after 26-week high fat diet^22^. *Cd68* and *Ftl* were co-expressed and markedly elevated in macrophage-like SMC clusters 1, 2 and 3, compared to other SMC clusters.

**Fig. S7. Putative regulators for macrophage-like SMCs. (A)** Ranked list of transcription factors (TFs) and regulators for SMC8-Mac according to degree centrality. **(B)** UMAP plot showing elevated expression of the 4 TFs in SMC8-Mac. **(C)** Network centrality analysis plot of TFs (*SPI1, MAFB, JUNB* and *KLF4*) revealing scores according to degree centrality, betweenness centrality and eigenvector centrality, highlighting their particular and consistent high scores for SMC8-Mac.

**Fig. S8. *In silico* knockout of putative master regulatory TFs in macrophage-like SMCs reverses phenotypic switching and shift towards contractile state. (A)** The effect of *in silico* knockout of *SPI1, MAFB, JUNB* and *KLF4* with CellOracle on SMC clusters. **(B)** Trajectory inference with regulatory directionality integrating pseudotime and perturbation predictions. The inner product score is measured, which shows directionality after TF perturbation, overlaid over a pseudotime-aligned UMAP. Green depicts cells that align positively with the TF-predicting direction. Purple depicts cells that oppose the TF direction after perturbation. The SMC8-Mac population (selected in orange) showed the strongest directional change after perturbation among all SMC clusters. **(C)** Gene expression of *FTL* in primary aortic smooth muscle cells after experimental *in vitro* knockdown of TF (*SPI1, KLF4, ATF3, MAFB, MYC, JUNB*) using siRNA. Data is represented as mean±SEM. (n=3 independent experiments); Student’s t-test.

**Fig. S9. RNA-seq analysis of macrophage-like switching in aortic SMCs after oxLDL stimulation. (A)** Principal component analysis plot showing 4 treatment groups (n=3 each). **(B)** Heatmap showing DEGs comparing between the 4 treatment groups (n=3 each). A, arrested control; O, 50 µg/ml oxLDL; IPO (10 ng/ml IL-1β+10 ng/ml PDGF-BB+50 ug/ml oxLDL) and IPCO (10 ng/ml Il-1β + 10 ng/ml PDGF-BB + 10 µM CML + 50 ug/ml oxLDL). **(C)** Volcano plot showing DEGs of oxLDL treatment compared to arrested control. **(D)** Volcano plot showing DEGs of IPO treatment compared to arrested control. **(E)** MA plot showing DEGs of IPCO treatment compared to arrested control.

**Fig. S10. *In vitro* experiments showing iron promotes proliferation, macrophage-like switching and cell death. (A)** Quantification of primary human aortic SMCs proliferation after treatment for 96 h with either serum free arrest media or growth media with 2% FBS alone, with or without 30 µM FAS, 5 µM Fer-1, or 100 µM DFO. Data is shown as mean±S.E.M.; one-way ANOVA with Tukey’s multiple comparison test. **(B-C)** Viability of SMCs after treatment with serum free arrest media and 100 µM FAS, 300 µM FAS, 3 µM RSL, 10 µM RSL, 5 µM Fer-1, and/or 100 µM DFO after treatment for 72 h. Data is shown as mean±S.E.M. (n=3); one-way ANOVA with Tukey’s multiple comparison test. **(C-D)** Migration quantified by scratch assay of SMCs after 15 or 8 h treatment with serum free media or the addition of 0.1 µM RSL3, 100 µM FAS, 5 µM Fer-1 or 100 µM DFO. Data is shown as mean±S.E.M. (n=3); one-way ANOVA with Tukey’s multiple comparison test. Data is shown as mean±S.E.M. (n=3); one-way ANOVA with Dunnett’s multiple comparisons test. **(E)** Gene expression of *PLIN2* in SMCs quantified by RT-qPCR after 48 h stimulation with serum free arrest media or the addition of 25 µg/ml oxLDL, IPCO, OFAS, 5 µM Fer-1 and/or 100 µM DFO. Data is shown as mean±S.E.M. (n=3); one-way ANOVA with Tukey’s multiple comparison test. **(F)** Western blot of TFRC, CD68, FPN1, SPP1, LGALS3, FTL and GAPDH in SMCs after stimulation with 25 µg/ml oxLDL, 5 µM FAS, OFAS (oxLDL+FAS), 5 µM Fer-1 and/or 100 µM DFO for 48 h. **(G)** Ferrous iron (Fe^2+^) in primary human aortic SMCs visualized by FerroOrange staining, after stimulating with serum free media or addition of 10 µM FAS, 5 µM Fer-1 or 100 µM DFO for 24 h. Data is shown as mean±S.E.M. (n=3); one-way ANOVA with Tukey’s multiple comparison test.

**Fig. S11. Mouse carotid tissues.** Elastin staining of *Ldlr*^-/-^ mouse carotid arteries using Elastic Van Gieson staining, after high fat diet with and without wire injury under treatment with either vehicle, RSL, or Fer-1 administration, for 2 weeks post-injury. Scale bar, 300 µm.
